# A map of RBP-miRNA regulatory connections for deciphering the mechanisms of transcriptome remodeling in cancer

**DOI:** 10.64898/2026.09.29.755492

**Authors:** Gabrielle Perron, Mohan Amaravadi, Hsin-Wei Tseng, Pouria Jandaghi, Elham Moslemi, Tamiko Nishimura, Maryam Rajaee, Rached Alkallas, Bruce Culbertson, Kristle Garcia, Ian R. Watson, Hani Goodarzi, Thomas Duchaine, Yasser Riazalhosseini, Hamed S. Najafabadi

**Author notes:** Correspondence: HSN and YR.

## Abstract

Micro-RNAs (miRNAs) are key post-transcriptional regulators of gene expression that influence cancer progression and outcome. Whereas transcriptional dysregulation of miRNA abundance underlies abnormal functionality of some miRNAs in cancer, emerging data points to RNA-binding proteins (RBPs) as prominent post-transcriptional regulators of miRNA activity. Here, by modelling the regulon activities of well-characterized miRNA families in RNA-seq data of a comprehensive collection of RBP inhibition experiments, human cell lines and tissues, we report a map of regulatory connections between 34 RBPs and 28 miRNAs. Furthermore, through a global analysis of the abundance as well as the activity of miRNAs in thousands of tumor and control samples from 17 human cancers, we show that RBP-driven regulation of miRNA activity is common in tumors and contributes to the heterogeneity in miRNA activity profiles. We uncover a set of RBPs that regulate the activity of tumor-suppressor and oncogenic miRNAs, including miR-29-3p, miR-17-5p and miR-19-3p. Of particular interest, we describe a pan-cancer negative regulatory role for DDX55 on the activity of the miR-29-3p family, a critical miRNA family with tumor suppressor functions that is involved in most human cancers. Our functional studies confirm that DDX55 inhibition restores the gene-silencing activity of miR-29-3p and reduces cancer cell viability. Overall, our study provides the first pan-cancer investigation of functional regulatory connections between RBPs and miRNAs, and highlights RBPs whose interventional modulation may serve as a new approach to reprogram abnormal miRNA activities in cancer.

## INTRODUCTION

Cancer is characterized by widespread transcriptome aberrations that affect thousands of genes and a variety of molecular pathways and cellular processes ^1,2^. For decades, post-transcriptional regulatory mechanisms have been acknowledged as playing key roles in cancer-associated gene dysregulation. Specifically, a large body of literature has highlighted different microRNAs (miRNAs) whose abnormalities act as master regulators of transcriptome remodeling in cancer. miRNAs are small non-coding RNAs that collectively regulate the expression of most protein-coding genes in the human genome ^3^. In general, miRNAs mediate RNA silencing, with miRNA-mRNA interactions leading to translational repression and/or destabilization of the mRNA target. miRNA dysregulation has been observed in most human cancers and has a causal role in tumorigenesis, with implications in virtually all hallmarks of cancer development and disease progression ^4^. Accordingly, miRNAs have been considered as potential biomarkers for diagnostic and prognostic purposes, or as potential therapeutic targets against cancer cells ^5^.

Similar to miRNAs, RNA-binding proteins (RBPs) are prominent post-transcriptional regulators of gene expression that modulate the stability or splicing of their target mRNAs through sequence-specific binding or recognition of structural and contextual features ^6,7^. Furthermore, RBPs are involved in virtually all aspects of miRNA biogenesis, turnover and loading of the RNA-induced silencing complex (RISC), thereby regulating miRNA-mediated gene silencing (miRNA activity) ^8,9^. As such, abnormal expression of RBPs has been shown to play important roles in shaping cancer transcriptomes ^10–12^. While recent studies have attempted to systematically identify RBPs that directly bind to miRNAs ^13,14^ or affect RNA secondary structures in the vicinity of miRNA-binding site ^15^, it is not clear to what extent such interactions may lead to functional changes in miRNA activity. In this study, we present a comprehensive map of direct and indirect RBP-miRNA regulatory axes, consisting of RBPs whose modulation leads to downstream alteration of the activity of specific miRNA(s). We find that for most RBP-miRNA regulatory pairs, RBP expression is associated with miRNA activity across a variety of cells and tissues. Through functional experiments, we confirm that inferred RBP-miRNA connections are reproducible and not limited to specific cell types. In addition, using data from thousands of cancer samples from the Cancer Network Atlas (TCGA), we show that RBPs regulate the activity of specific tumor suppressor and oncogenic miRNAs, thereby impacting the phenotypic behaviour of cancer cells. Specifically, we show that upregulation of DDX55, which is associated with poor prognosis, inhibits the activity of the tumor suppressor miR-29-3p family across multiple cancer types and that DDX55 inhibition reduces the viability of cancer cells. These results provide insights into the mechanisms of miRNA dysregulation and transcriptome remodelling in cancer.

## RESULTS

### RBPs regulate miRNA-mediated stability programs

To explore functional connections between RBPs and miRNAs, we first modelled the functional status (activity) of each miRNA using a scalable framework that considers the stability of the miRNA target mRNAs, its regulon, as a readout of miRNA activity (**Fig. 1**). This framework is based on our recent computational methods for inference of mRNA stability by joint analysis of exonic and intronic reads in RNA-seq data ^16^ and network modeling of regulon activities ^17^ (see Methods). The network of binding interactions between miRNAs and target mRNAs (predicted miRNA regulons) was built based on matches to the seed sequence of 153 conserved miRNA families (**Supplementary Table 2**). We examined the ability of this network for capturing valid miRNA-mRNA interactions by performing AGO2 cross-linking ligation and sequencing of hybrids (AGO2-CLASH) ^18^ for 14 predicted interactions (**Supplementary Fig. 1a**). The AGO2-CLASH experiments confirmed 50% of the tested interactions, whereas it did not detect any hybrids for 9 control miRNA-mRNA pairs without matched seeds (**Supplementary Fig. 1b**). These results demonstrated the ability of the network to capture information about true miRNA-mRNA interactions, with a positive predictive value of 50%. Using this constructed network, we interrogated miRNA activity changes in 468 publicly available RNA-seq datasets of shRNA-medicated RBP knockdown (KD) experiments, encompassing 230 unique RBPs, in human cell lines (HepG2 and K562) from the Encyclopedia of DNA elements (ENCODE) ^19^ (**Supplementary Table 1**). Briefly, in each sample, we obtained batch-corrected genome-wide profiles of mRNA stability changes relative to the average across experiments (See **Supplementary Fig. 2 and 3**, and Methods). Then, we modeled these changes occurring upon each RBP perturbation as a function of the destabilizing effects of miRNAs that bind to the 3’UTR of each mRNA, enabling us to infer the differential miRNA activity in each sample (**Fig. 1**). We also corrected for shRNA-mediated off-target effects, which appeared to stem from the presence of a partial match between shRNA helices and 3’ UTRs (**Supplementary Fig. 4 and 5, Supplementary Table 1**). Subsequently, we identified 39 regulatory interactions between 34 RBPs and 28 miRNAs, which were statistically significant in at least one biological replicate (FDR < 0.05), with other replicates showing consistent effect (**Fig. 2a-b; Supplementary Table 3**). Noticeably, one network of interconnected RBPs and miRNAs stood out, involving 9 RBPs (DDX55, PTBP1, SUPV3L1, MSI2, RPLP0, FIP1L1, FAM120A, IGF2BP2 and PABPC4) and 4 miRNA families (miR-29-3p, miR-19-3p, miR-218-5p and miR-17-5p^1^). These results suggest that a given RBP may modulate the activity of specific sets of miRNAs.

**Figure 1.**
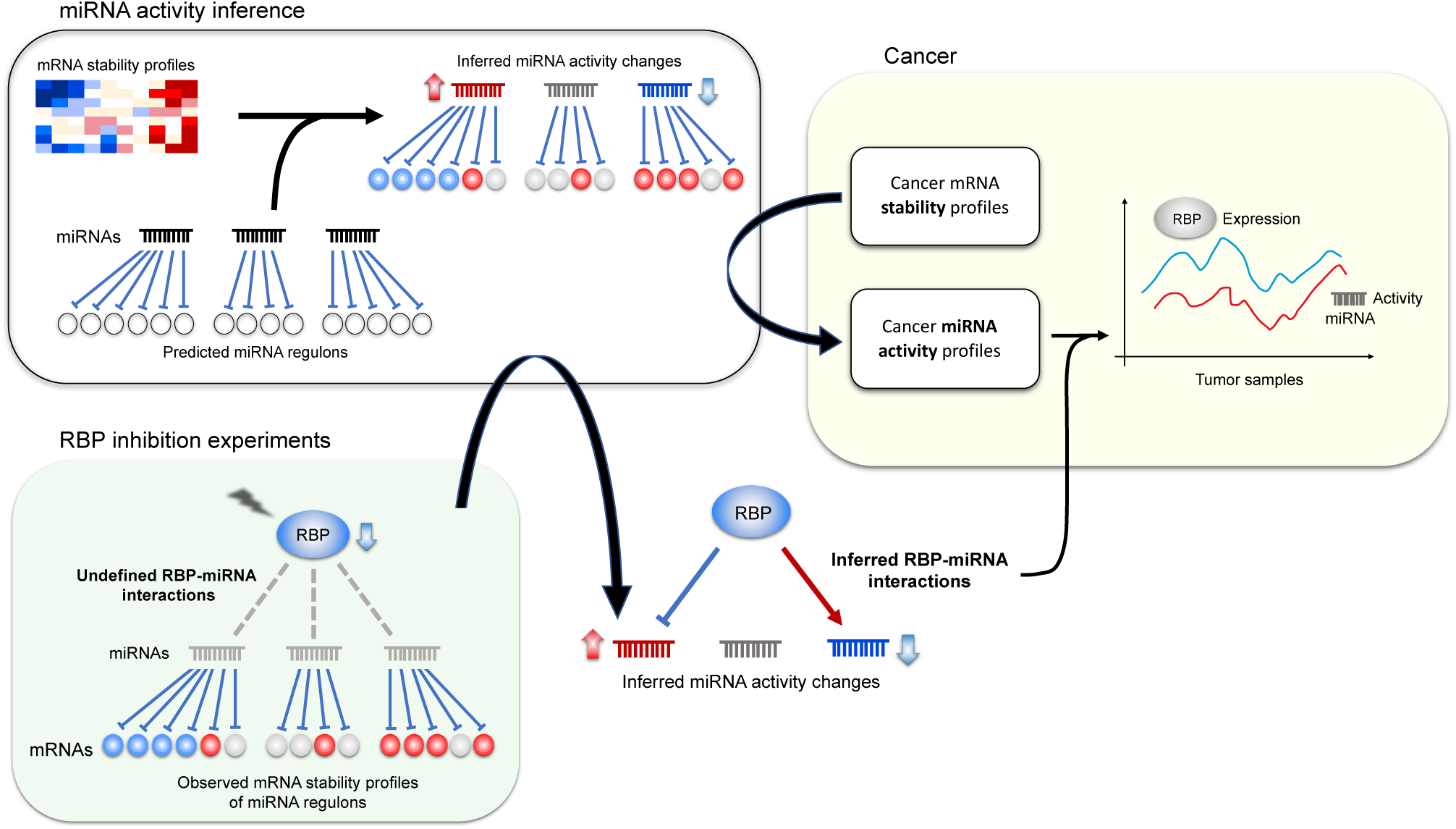
Schematic representation of the inference of miRNA activities, functional interactions between RBPs and miRNAs and their relevance to cancer. mRNA stability profiles derived from RNA-seq data of RBP knockdown experiments were combined with predicted miRNA regulatory networks to infer changes in miRNA activity induced by RBP inhibition. Functional relationships between RBPs and miRNAs were then deduced from the change in miRNA activity upon inhibition of each RBP. See Methods for more details. This framework was then applied to RNA-seq data of TCGA samples to infer miRNA activity changes in tumors. The relevance of the regulatory connections between RBPs and miRNAs was assessed in the context of cancer, notably by investigating the association between RBP expression and miRNA activity across various cancer types.

**Figure 2.**
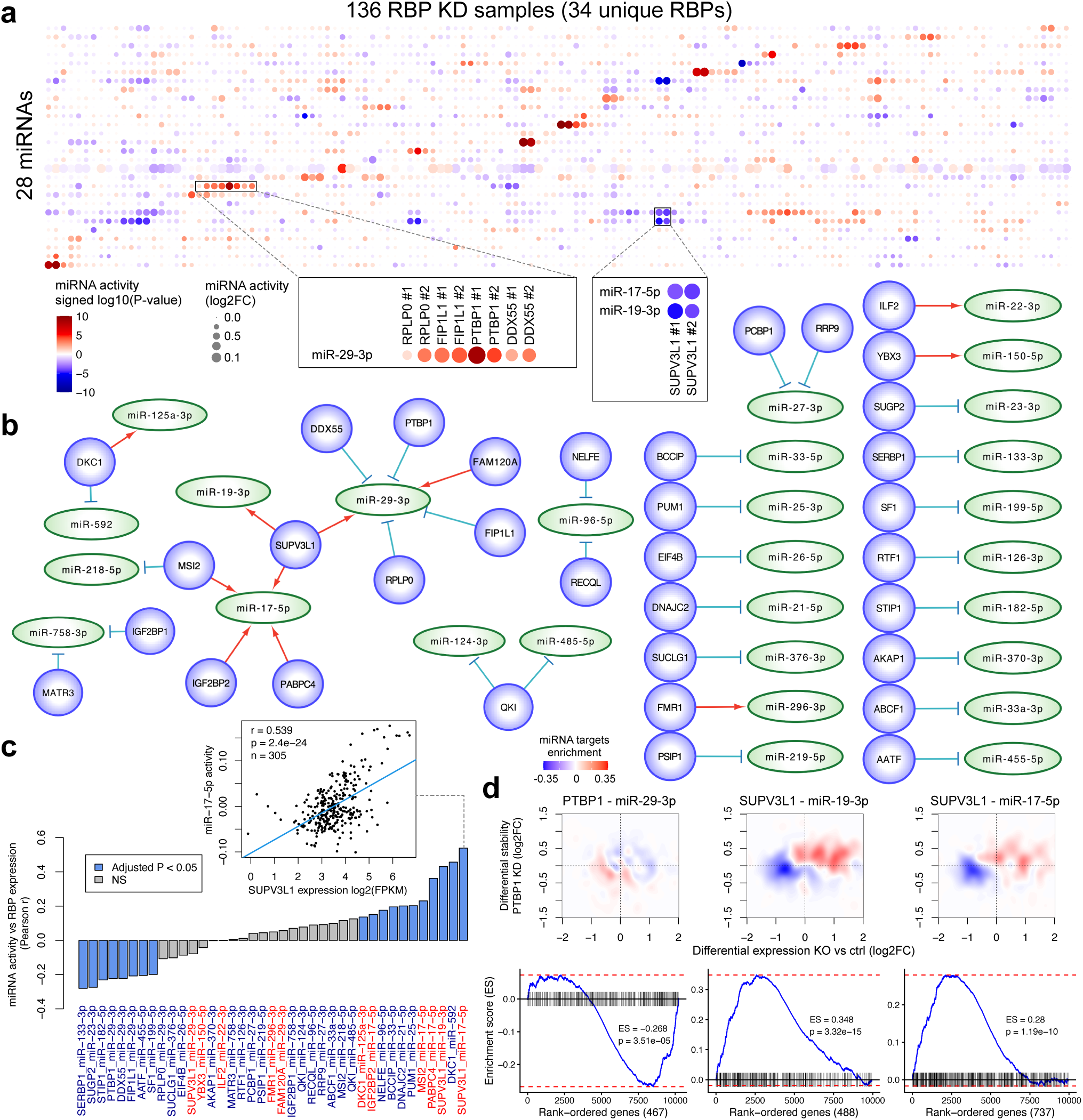
RBP inhibition reproducibly affects miRNA activity. (**a**) Inferred miRNA activity profiles in systematic RBP shRNA-mediated knockdown experiments in HepG2 and K562 cells from ENCODE, following correction of mRNA stability for shRNA off-target effects. The color gradient represents signed log10(P-values) associated with inferred differential miRNA activity, with red indicating increased miRNA activity relative to the average, and blue decreased miRNA activity. The dot size represents the log2 fold-change of differential miRNA activity. miRNAs and knockdown samples with at least one significant result at FDR < 0.05 were retained, along with all their biological replicates in both cell lines. A few example RBP-miRNA connections of interest to this study are highlighted. (**b**) Networks of significant RBP-miRNA connections for which biological replicates were in agreement in terms of type of interaction (activating or inhibitory). The edges represent the type of interaction, with a red arrow representing an activating interaction, and a blue blunt arrow representing an inhibitory interaction. These interactions are listed in **Supplementary Table 3**. Networks were drawn using Cytoscape 3.9.1 ^56^. (**c**) Pearson correlations of inferred miRNA activity and RBP expression (log2 FPKM) for the candidate RBP-miRNA regulatory pairs, across external RNA-seq datasets from ENCODE including a variety of cell lines and tissue types. Blue bars represent significant associations at FDR < 0.05. The font color of the x-axis labels represents the type of inferred interaction, with red indicating an activating interaction, and blue an inhibitory interaction. The scatter plot shows the association between the expression of SUPV3L1 and the activity of miR-17-5p, as an example. (**d**) The plots at the top show the enrichment of miRNA targets among differentially stabilized genes in RBP knockdown, compared to knockout experiments, for three candidate interactions involving the RBPs PTBP1 and SUPV3L1. RBP knockdown measurements are from the cell line in which the interaction was significant (see **Supplementary Table 3**). The x-axis represents the log2 fold-change of differential expression in the RBP knockout relative to the empty vector controls, while the y-axis represents the median log2 fold-change of mRNA stability in the RBP knockdown, relative to the average. The color gradient shows the difference between the density of genes that are miRNA stability targets and that of other genes. Red represents a higher density of miRNA targets compared to other genes, while blue represents a lower density. The plots at the bottom show gene set enrichment analyses (GSEA^57^; using the ‘fgsea’ R package version 1.32.0) for RBP knockouts, using the Wald statistic of differential expression as the ranking metric. The blue line represents the running enrichment score (ES) for RBP-responsive miRNA targets (black lines), for which the differential stability was in the expected direction in the RBP knockdowns, across the rank-ordered gene list (of the RBP knockouts; x-axis).

**Figure 3.**
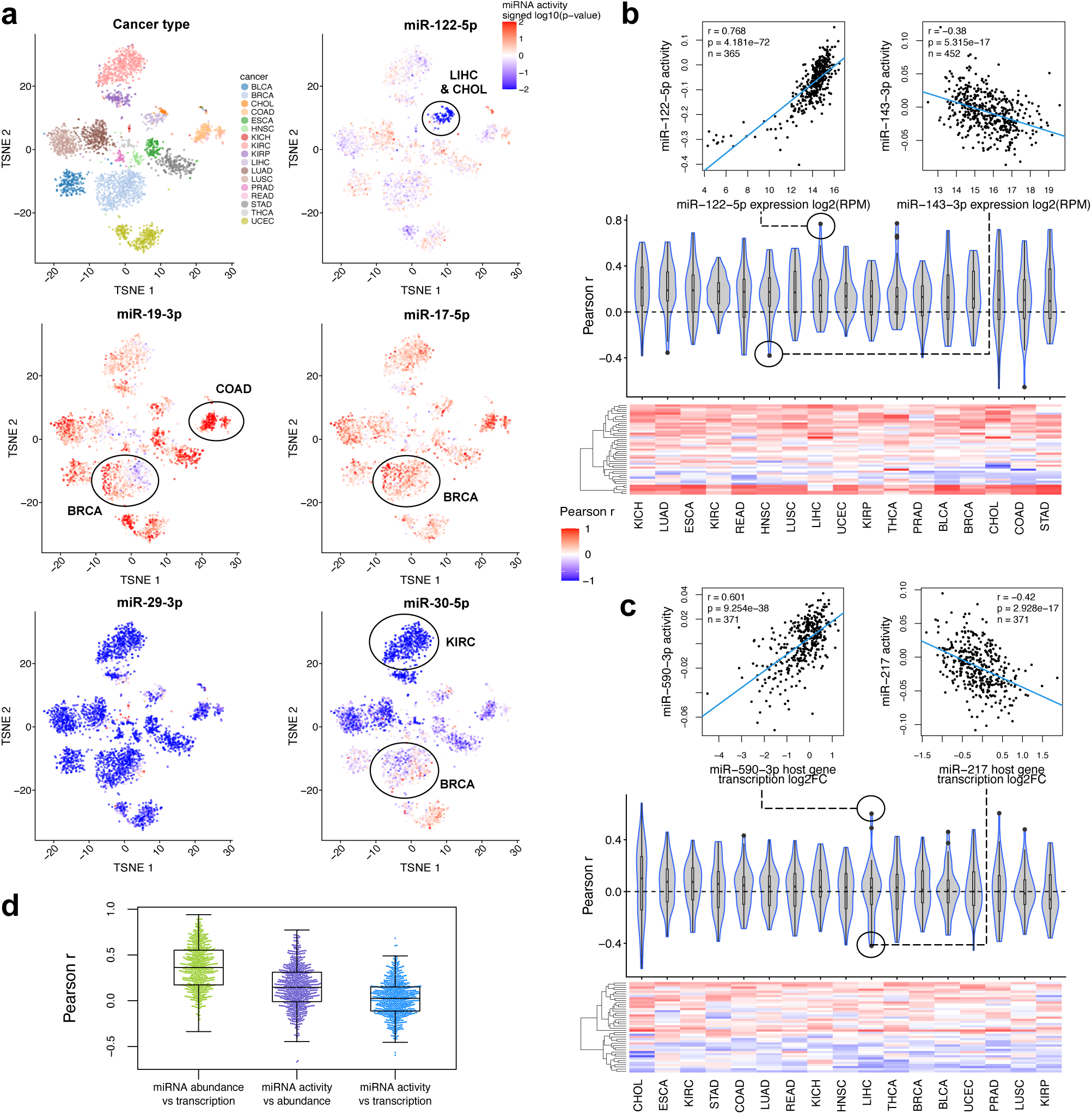
Pan-cancer miRNA activity profiles and contributions of miRNA transcription and abundance. (**a**) t-SNE plots showing inferred differential miRNA activity profiles in TCGA tumors. Each point represents a tumor sample, and the colors represent the cancer types in the first plot (top-left), and differential miRNA activity, relative to the average of normal samples, for 5 different miRNAs in the subsequent plots. The color gradient represents the signed log10(P-value) associated with inferred differential miRNA activity, with red indicating increased miRNA activity in tumors relative to normal tissues, and blue decreased miRNA activity. Dimensionality reduction was performed on the miRNA activity data for 3162 tumors with at least one significant miRNA, and 91 miRNAs that were significant in at least one tumor. t-SNE plots were generated using the ‘scater’ R package ^58^. (**b**) Association between the activity of each miRNA family and the abundance of each corresponding mature miRNA family member in TCGA tumors, for miRNA families with significant changes in activity in at least 10 tumor samples in at least one cancer type (see **Supplementary Table 5**). Both the violin plots and heatmap show the Pearson correlation coefficients for each cancer type. Each data point in the violin plots and each row in the heatmap represent one mature miRNA. The scatter plots show the association between miRNA activity and abundance across tumors for two example miRNAs in two different cancer types. (**c**) Similar to (b), for the association between miRNA activity and miRNA transcription (instead of abundance) in TCGA tumors. Transcriptional changes were inferred from that of the host genes, based on changes in intronic read counts (see Methods). (**d**) Box plot comparing the Pearson correlation coefficients for the associations between miRNA abundance vs transcription, (see **Supplementary Fig. 9b**), activity vs abundance (see b), and activity vs transcription (see c).

To examine the reproducibility of the identified RBP-miRNA interactions, we interrogated the association between RBP expression and miRNA activity for each pair in an independent set of 308 RNA-seq profiles from the ENCODE database, which include samples from tissues, primary and immortalized cell lines (**Supplementary Table 1**). This analysis showed correlations between RBP expression and miRNA activity in agreement with shRNA-mediated inhibition for 20 out of the 39 RBP-miRNA interactions that we had inferred (**Fig. 2c**), which we considered as a set of high-confidence interactions. The concordance increased with the stringency of the association testing; for example, out of 14 RBP-miRNA pairs with significant correlation (FDR < 0.05) and |r| > 0.2, 11 were in the expected direction.

To validate the robustness of our computationally inferred RBP-miRNA regulatory connections using an orthogonal approach, we analyzed the transcriptional profiles of HEK293T cells following CRISPR-mediated knockout of three RBPs implicated in our RBP-miRNA interaction network: DDX55, PTBP1 and SUPV3L1 (**Supplementary Table 1**). While viable SUPV3L1 knockout clones could not be obtained, we were able to retain one clone with reduced expression (**Supplementary Fig. 6a**). Obtaining DDX55 knockout proved especially challenging despite multiple attempts, as DDX55 appeared to be essential for cell survival, with null expression being lethal. In cases where clones with mutations causing reduced expression of DDX55 were viable, the full length protein was still expressed and mutations were often in-frame, indicating that the protein may have been functional (**Supplementary Fig. 6b**). Furthermore, we observed substantial variability in the expression data between these samples (**Supplementary Fig. 6d**). We therefore excluded DDX55 from this analysis and instead further interrogated the DDX55-miR-29 connection using other experimental approaches, which are discussed later on in the study. Complete PTBP1 knockouts were successfully generated (**Supplementary Fig. 6c**). In total, we retained three PTBP1 knockout clones and one clone with reduced SUPV3L1 expression (see Methods and **Supplementary Fig. 6**). Using RNA-seq, we confirmed that transcriptional changes induced by these CRISPR-mediated RBP knockouts were consistent with those observed upon shRNA-mediated inhibition (**Fig. 2d**). More specifically, miR-29-3p targets were enriched among genes that were downregulated upon PTBP1 knockout, consistent with the inferred negative regulatory role of PTBP1 on miR-29-3p activity. Similarly, miR-17-5p and miR-19-3p targets were enriched among genes that were upregulated after SUPV3L1 knockout, in agreement with the ENCODE data showing reduced activities of these miRNAs upon SUPV3L1 inhibition. Taken together, these results support the RBP-miRNA regulatory connections inferred by applying our framework to the RBP knockdown data from ENCODE and show that many of these interactions are not cell type or assay-specific. Therefore, we sought to apply this framework to cancer transcriptomes to identify RBP-miRNA regulatory interactions that are relevant to cancer.

### Dysregulation of miRNA transcription and abundance only partially explains cancer-associated miRNA activity profiles

To generate pan-cancer miRNA activity profiles, we applied our miRNA activity inference framework to RNA-seq data from 6903 tumors and 677 adjacent normal samples across 17 solid tumor types from TCGA. Briefly, we obtained genome-wide mRNA stability changes relative to the average of cancer type-matched healthy tissues and inferred the differential activity of 153 miRNA families in individual tumor samples (**Fig. 1, Supplementary Table 2, Supplementary Table 1**; see Methods). We observed changes in the activity of a wide range of miRNAs in cancer; overall 91 miRNAs were modulated in at least one tumor sample relative to normal tissues (FDR < 0.05) (**Supplementary Fig. 7 and Supplementary Table 6**). Several miRNAs appeared to be recurrently dysregulated in a cancer-specific manner (**Fig. 3a, Supplementary Fig. 7 and Supplementary Table 5**). For example, 20% of colon adenocarcinomas (COAD) show a significant increase in miR-19-3p activity, consistent with the previously demonstrated oncogenic role of miR-19-3p in this cancer ^20,21^. Likewise, 83% of clear cell renal cell carcinomas (KIRC) show decreased activity of miR-30-5p, in line with previous literature on its tumor suppressive function and potential as a biomarker in KIRC ^22–24^. miR-122-5p, the miRNA that is most abundantly expressed in human liver cells ^25^, shows decreased activity in 16% and 97% of liver hepatocellular carcinoma (LIHC) and cholangiocarcinoma (CHOL) tumors, respectively, corroborating previous literature ^26,27^. In addition, we observed consistent pan-cancer dysregulation for several well-established tumor suppressor or oncogenic miRNAs (**Fig. 3a, Supplementary Fig. 7, Supplementary Table 5**). Notably, miR-29-3p activity appeared reduced in the majority of tumor samples across several cancer types, in agreement with its known tumor suppressive function and general downregulation in most human cancers ^28^. Numerous samples from different cancers were also characterized by increased miR-19-3p and miR-17-5p activity. These miRNAs are part of the oncogenic miR-17∼92 cluster, also known as oncomiR-1, which has a well-established oncogenic function and shown to be amplified in different cancers ^29,30^. We also observed substantial heterogeneity in miRNA activity within most cancer types, especially in the case of miR-19-3p, miR-17-5p and miR-30-5p, including in breast invasive carcinoma (BRCA) (**Fig. 3a and Supplementary Fig. 7**).

We began to investigate the underlying mechanisms of this heterogeneity in miRNA activity between TCGA tumors by evaluating the correlation between inferred miRNA activity profiles and mature miRNA abundance measurements from small RNA-seq (**Supplementary Table 1**). Within each cancer type, an average of 75% of miRNAs families showing significant tumor-associated activity changes displayed a positive correlation (not considering the significance) between activity and abundance (compared to only 51% for non-matching pairs of miRNA families and mature miRNAs; **Fig. 3b and Supplementary Fig. 8**). The overall positive association between inferred activities and sequencing-based miRNA abundances confirms that cancer-associated changes in abundance partially explain the differential activity patterns observed for most miRNAs. We note, however, that the strength of the associations are still relatively weak (average Pearson *r*=0.15). In addition, miRNA activity is even more weakly associated with the rate of transcription of their primary transcripts, estimated by quantifying the changes in intronic reads of miRNA host genes (average *r*=0.024; **Fig. 3c-d, Supplementary Fig. 9a and Supplementary Table 1**; see Methods). Furthermore, even when selecting for differentially expressed miRNAs, abundance itself is only partially explained by the rate of transcription of their primary transcripts (average *r*=0.37; **Fig. 3d and Supplementary Fig. 9b**), although this association is expectedly stronger than that between miRNA abundance and activity, or between miRNA activity and host gene transcription. Indeed, while host gene transcription almost always positively correlates with mature miRNA abundance within each tumor type, on average it accounts for only ∼19% of the variance in miRNA abundance. These observations suggest that, despite widespread variations in miRNA activity between tumors of the same cancer type, miRNA host gene transcription only partially explains the mature miRNA abundance, and mature miRNA abundance only partially explains the miRNA regulon activity, highlighting the involvement of substantial post-transcriptional modulation of miRNA activity in cancer.

### RBPs are key regulators of miRNA activity profiles in cancer

Given our preceding findings that RBPs regulate miRNA activity patterns in cell lines and tissue samples, and that miRNA transcription and abundance do not fully explain miRNA activity profiles in cancer, we investigated the role of RBPs in miRNA regulation in tumors. We first noticed that changes in RBP expression between tumor and normal samples generally appeared to mirror cancer-associated miRNA activity patterns (**Fig. 4a**). For example, DDX55 and PTBP1 are consistently upregulated in tumors, which is coherent with a global downregulation in the activity of miR-29-3p, the miRNA that these RBPs were inferred to inhibit (**Fig. 2**). In the case of SUPV3L1, its expression is overall increased in tumors, but with substantial heterogeneity within and between cancer types. Interestingly, these heterogeneous changes are mirrored by the changes in miR-17-5p and miR-19-3p activities and are consistent with the inferred activating effect of SUPV3L1 on these miRNAs (**Fig. 4a**). That is, tumors with SUPV3L1 upregulation display an increase in miR-19-3p and miR-17-5p activities, and vice-versa for tumors with reduced SUPV3L1 expression. We systematically examined the correlations between miRNA activity and RBP expression levels within each TCGA cancer types for the 20 high-confidence RBP-miRNA pairs (**Fig. 2c**). Of the 19 interactions for which expression data for the RBP was available, 12 showed correlation patterns consistent with the predictions in at least 70% of the studied TCGA cancer types (**Fig. 4b**). These associations were maintained after correcting for tumor purity, which is also associated with miRNA activity (**Supplementary Fig. 10a-b**; see Methods). Together, these results show that within each cancer type, associations between miRNA activity profiles and the expression of their predicted RBP regulators are consistent with the inferred regulatory direction (activating or inhibitory), highlighting the potential role of RBPs in tumor-specific dysregulation of miRNA activity.

**Figure 4.**
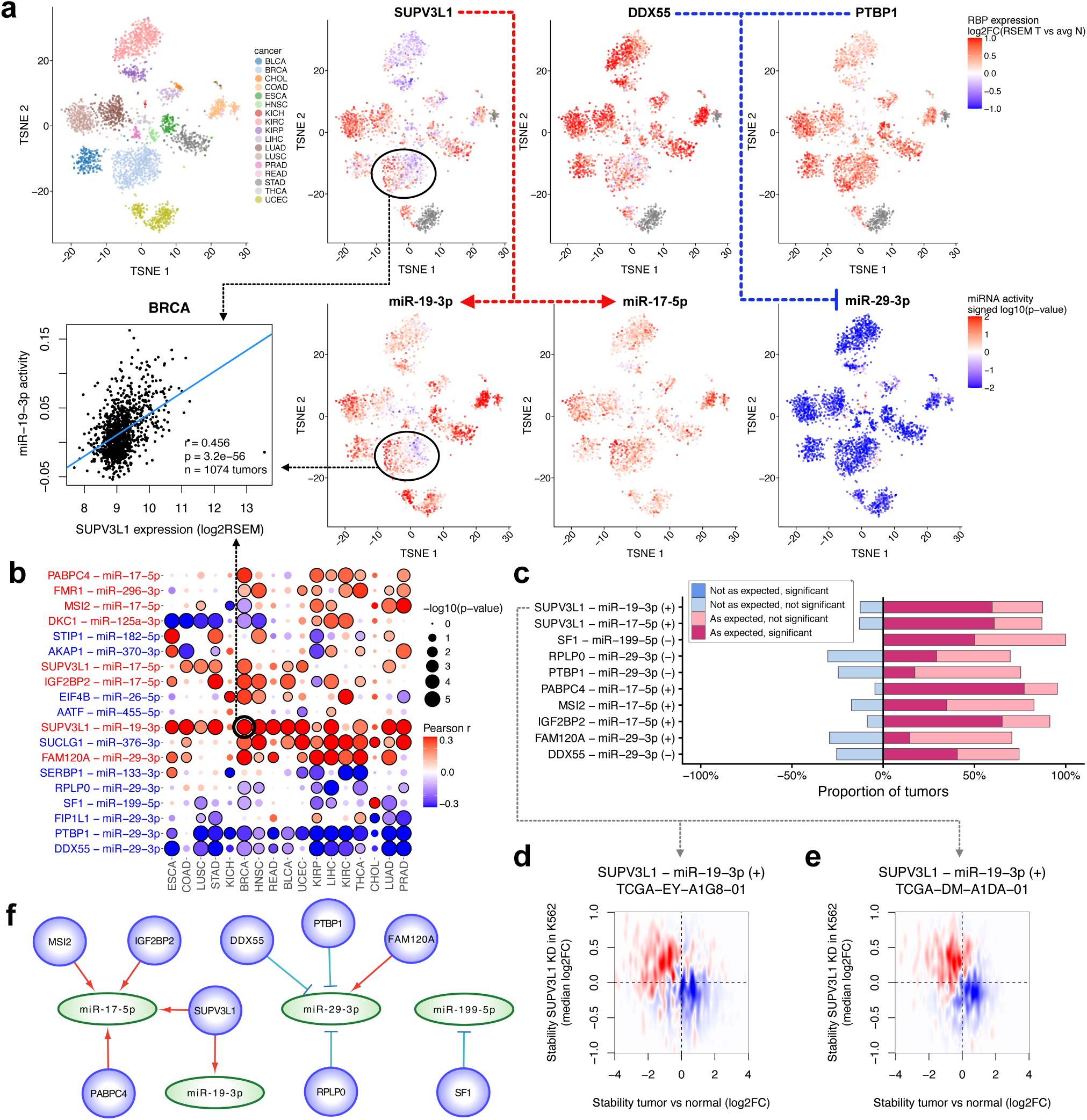
Pan-cancer relevance of RBP-miRNA connections. (**a**) t-SNE plots showing inferred miRNA activity and RBP expression profiles in TCGA tumors, for four example RBP-miRNA regulatory pairs involving 3 miRNAs (miR-29-3p, miR-19-3p and miR-17-5p) and 3 RBPs (DDX55, PTBP1 and SUPV3L1). Each point represents a tumor sample. The color gradients represent the differential miRNA activity for the miRNA plots, and the gene expression log2(RSEM) fold-change in tumors for RBP plots, both relative to the average of normal samples. Dimensionality reduction was performed on the miRNA activity data retaining samples and miRNAs with at least one significant hit at FDR < 0.05. The arrows represent the type of interaction, where a red arrow represents an activating interaction, and a blue blunt arrow represents an inhibitory interaction. t-SNE plots were generated using the ‘scater’ R package ^58^. The scatter plot shows the association between SUPV3L1 expression and miR-19-3p activity in BRCA as an example. (**b**) Pearson correlations between inferred miRNA activity and RBP expression across tumors, for each high-confidence regulatory pair (for which the correlations in ENCODE cell lines and tissues were in the direction expected from the interaction type inferred from the RBP KD experiments; see Fig. 2c). The circle size represents the -log10(P-value) of correlations, while the color gradient represents the correlation coefficients. A black outline indicates a significant result at FDR < 0.05. The font color of the row labels represents the type of inferred interaction, with red indicating an activating interaction, and blue an inhibitory interaction. (**c**) For each candidate RBP-miRNA interaction from (b) that had correlations consistent with the predicted type of interaction in at least 70% of TCGA cancers (y-axis, with the inferred type of the interaction, activating “+” or inhibitory “-“, in parentheses), the predicted targets of the miRNA that responded to RBP knockdown in vitro in the direction consistent with that interaction type were tested for enrichment among stabilized or destabilized mRNAs in every TCGA tumor sample showing a significant (FDR < 0.05) change in miRNA activity (tumor vs. normal), using GSEA^57^ (‘fgsea’ R package version 1.32.0) and using mRNA stability estimates relative to the average of normal samples as the ranking metric. Bars show, per interaction, the proportion of tested tumors with RBP-KD-consistent predicted miRNA targets enriched as expected or not (based on the change in miRNA activity in the sample and on the type of interaction), split into significant (FDR < 0.05) and non-significant fractions. (**d**) Enrichment of miR-19-3p targets among differentially stabilized genes in the SUPV3L1 knockdown experiment in K562 cells and in the TCGA-UCEC (Uterine corpus endometrial carcinoma) tumor TCGA-EY-A1G8-01, as an example. The x-axis represents the log2 fold-change of stability in tumor relative to the average of normal samples, while the y-axis represents the median log2 fold-change of stability in the SUPV3L1 knockdown experiments relative to the average. RBP knockdown data are from the cell line where the interaction was inferred (see **Supplementary Table 3**). The color gradient shows the difference between the density of genes that are miRNA stability targets and that of other genes. Red represents a higher density of miRNA targets compared to other genes, while blue represents a lower density. (**e**) Similar to (d), here showing, as a second example, the enrichment of miR-19-3p targets among differentially stabilized genes in the SUPV3L1 knockdown experiment in K562 cells, and in the TCGA-COAD tumor TCGA-DM-A1DA-01. (**f**) Networks of RBP-miRNA connections with pan-cancer relevance. These represent regulatory connections from (b) that were consistent with the predictions in terms of correlation between RBP expression and miRNA activity in at least 70% of TCGA cancers, and for which the same miRNA targets appear to be affected in the expected direction upon RBP inhibition and in cancer in the majority of tumors, regardless of significance, in (c). The edges represent the type of interaction, where a red arrow represents an activating interaction, and a blue blunt arrow represents an inhibitory interaction. These interactions are listed in **Supplementary Table 6**.

To further confirm the regulatory roles of the identified RBPs on corresponding miRNAs in cancer, we asked if the same predicted miRNA targets that were responsive to *in vitro* RBP inhibition are also dysregulated in TCGA tumors. We examined whether the cancer-associated (tumor vs. normal) stability profiles of these specific targets were in line with the inferred RBP-miRNA regulatory connections. Overall, for the 10 interactions where the miRNA shows a significant change in activity in at least one tumor, the miRNA targets responsive to *in vitro* RBP inhibition were overall dysregulated in the expected direction in at least 70% of tumor samples where the miRNA was differentially active, regardless of significance (**Fig. 4c**). For example, we observed an enrichment of miR-19-3p target mRNAs that were stabilized upon SUPV3L1 knockdown, among destabilized genes in tumors with increased miR-19-3p activity (**Fig. 4d-e**), in line with the inferred activating interaction for this pair (**Fig. 2b and Supplementary Table 3**). Notably, whereas significant enrichments (|ES| > 0 and FDR < 0.05) of expected sign were observed on average in 45% (up to 77%) of tumors, significant enrichments in the opposite direction were observed in almost none of the tumors (0.35%; **Fig. 4c**). Although these miRNA target enrichments were less significant when predicted based on RBP expression instead of miRNA activity, most of the significant results were still as expected (**Supplementary Fig. 11**). We therefore considered the aforementioned 10 RBP-miRNA connections as our final set of interactions with pan-cancer relevance (**Fig. 4f, Supplementary Table 6**). These networks highlight miR-29-3p and miR-17-5p as two miRNA families that are regulated by multiple RBPs, which ultimately impact the cancer transcriptome. Taken together, these results support the inferred RBP-miRNA connections and uncover an important role for RBPs in regulating the activity of key miRNAs in cancer.

### DDX55 inhibits the activity of miR-29-3p and is essential for tumor cell viability

Notably, our analysis revealed that several RBPs appear to modulate the known tumor-suppressive miR-29-3p family, which also showed the largest change in activity across the greatest number of cancer types in our pan-cancer analysis of miRNA activity (**Supplementary Fig. 7 and Supplementary Table 5**). Our modeling of RBP-miRNA networks identified DDX55, PTBP1 and RPLP0 as negative regulators and FAM120A as a positive regulator of miR-29-3p activity, their expression inversely correlated with miR-29-3p activity across cell line and tissue samples from ENCODE as well as in TCGA data (**Fig. 2, 4a-c, 4f, 5b, Supplementary Table 6, Supplementary Fig. 10b, 11 and 12b**). Among these RBPs, DDX55 and PTBP1 are also broadly upregulated in tumors (**Fig. 5a and Supplementary Fig. 12a**). Interestingly, a physical interaction between PTBP1 and miR-29 has been observed in a previous study, which demonstrated that PTBP1 directly binds to the miR-29b-2 and miR-29c pre-miRNA hairpins, further supporting our inferences ^13^. On the other hand, we did not find any previous reports on a relationship between DDX55 and miR-29-3p, prompting us to further characterize this inhibitory interaction. This regulatory effect is of particular interest since DDX55 expression is strongly associated with poor patient survival in multiple cancers (**Fig. 5c-d**), even after correcting for patient age, tumor stage and purity (**Supplementary Fig. 10c**). Given that DDX55 is inferred to inhibit miR-29-3p activity, this observation is consistent with the thoroughly documented role of miR-29-3p as a tumor suppressor and a marker of favorable prognosis ^28^.

**Figure 5.**
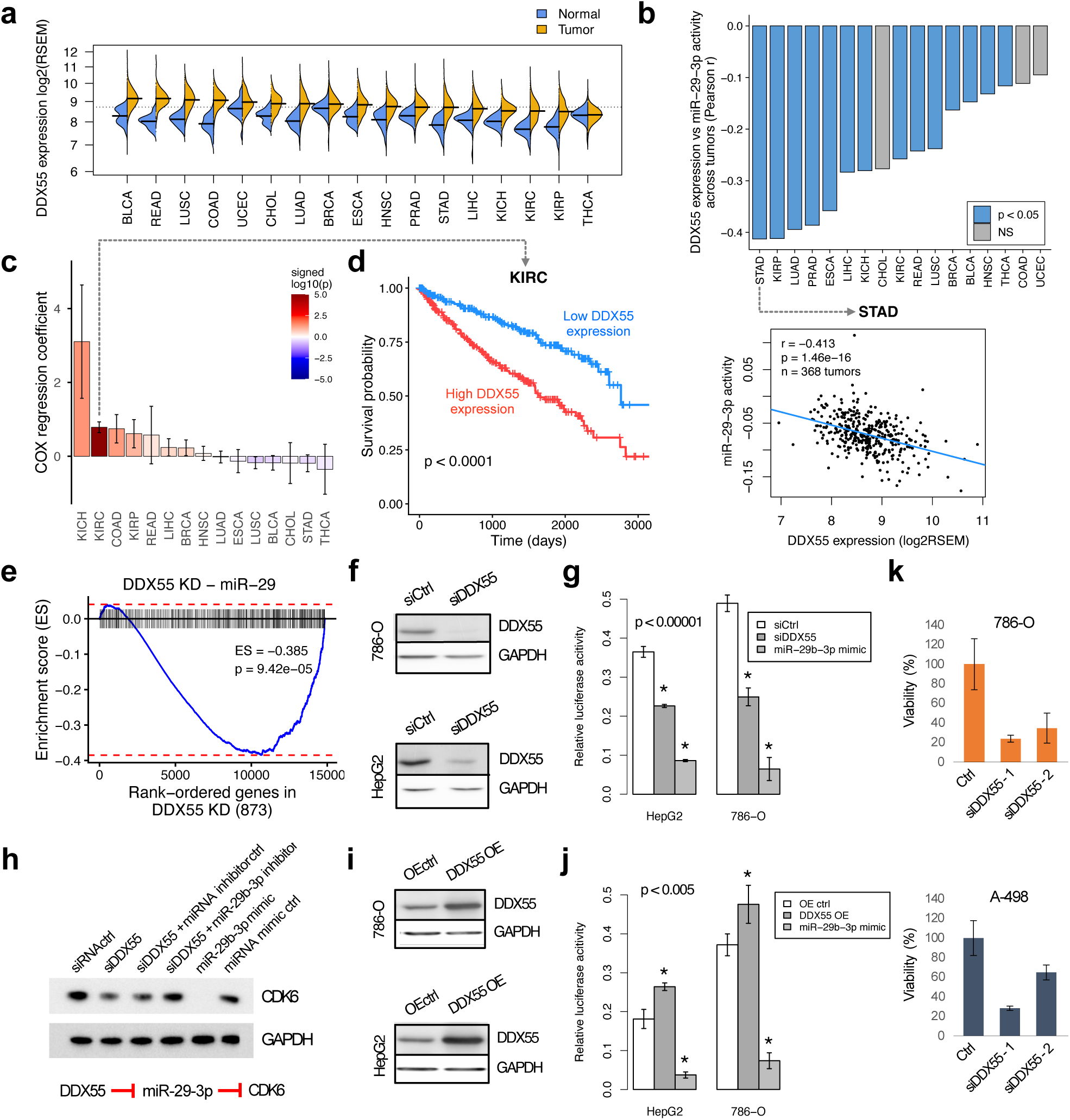
DDX55 inhibits the activity of miR-29-3p and is associated with patient survival and cell viability. (**a**) DDX55 expression across TCGA datasets, shown as log2(RSEM). (**b**) Bar plot showing the correlation between DDX55 expression and miR-29-3p activity across TCGA tumors. Blue represents significant associations (also see Fig. 4b). The scatter plot illustrates, as an example, the correlation between DDX55 expression and miR-29-3p activity in TCGA-STAD (Stomach adenocarcinoma), the cancer type where the association was most significant. (**c**) Association between DDX55 expression and patient survival, assessed using a Cox proportional hazards model adjusted for tumor stage. The color gradient represents signed log10(P-value). Red depicts an increased risk of death, while blue illustrates a decreased risk. Error bars represent SE. (**d**) Kaplan-Meier curve depicting the survival of TCGA-KIRC patients with high or low (above or below median) DDX55 expression. KIRC was selected as an example, as it showed the smallest P-value in (c). The P-value obtained using the log-rank test is included on the plot. (**e**) GSEA^57^ (‘fgsea’ R package version 1.32.0) for predicted miR-29 targets that were also either downregulated following miR-29 overexpression (mimic) in 786-O or A-498 cells, or upregulated following miR-29 inhibition in ACHN cells, in a DDX55 knockdown experiment from ENCODE in HepG2 cells. Genes (x-axis) are sorted by Wald test statistic of differential gene expression in the DDX55 knockdown experiment. The blue line shows the running enrichment score (ES) for miR-29 targets (black lines) across the rank-ordered gene list. (**f**) Western blot confirming the siRNA-mediated DDX55 knockdown in 786-O (top) and HepG2 (bottom) cell lines, compared to the non-targeting siRNA. (**g**) Relative luciferase luminescence (Renilla vs Firefly), representing the inverse of miR-29b-3p activity, upon DDX55 knockdown and subsequent expression of a miR-29b-3p mimic, or a siRNA negative control. Error bars represent SD, for 5 biological replicates. The significance of the change in relative luciferase activity relative to the control was determined using Student’s t-test. (**h**) Western blot showing the abundance of the miR-29-3p target CDK6 in 786-O cells expressing a control siRNA, an siRNA specifically targeting DDX55, the DDX55 siRNA and a miRNA inhibitor control, the DDX55 siRNA and a miR-29b-3p inhibitor, a miR-29b-3p mimic, and a miRNA mimic control. (**i**) Western blot confirming DDX55 overexpression in 786-O (top) and HepG2 (bottom) cell lines, compared to control. (**j**) Relative luciferase luminescence (Renilla vs Firefly), representing the inverse of miR-29b-3p activity, upon DDX55 overexpression and subsequent expression of a miR-29b-3p mimic, or an empty vector control (pXL5). Error bars represent SD, for 5 biological replicates. The significance of the change in relative luciferase activity relative to the control was determined using the Student’s t-test. (**k**) Viability of siRNA-mediated DDX55 knockdown 786-O and A-498 cell lines, as measured using the Sulforhodamine B (SRB) assay. The y-axis represents the % of viable cells. Error bars represent SD, for 5 biological replicates.

To experimentally validate the impact of DDX55 on functional miR-29 targets, we examined whether targets of miR-29 that are responsive to miR-29 over-expression and inhibition were also affected by DDX55 inhibition. We used data from a previous study where a miR-29b-3p mimic was expressed in A-498 and 786-O cell lines, and a miR-29b-3p inhibitor was expressed in ACHN cells ^31^. Predicted miR-29-3p targets that were downregulated when miR-29b-3p was overexpressed, or upregulated when miR-29b-30 was inhibited, were also enriched among destabilized genes upon DDX55 knockdown, consistent with an inhibitory interaction (**Fig. 5e**). To further validate the inhibitory effect of DDX55 on miR-29-3p, we inhibited DDX55 expression through siRNA-mediated knockdown in HepG2 and 786-O cancer cell lines (**Fig. 5f and Supplementary Fig. 13a**) and examined changes in miR-29-3p activity using dual luciferase reporter assays. miR-29b-3p activity was significantly increased in both cell lines upon DDX55 knockdown, as shown by reduced luciferase signal (**Fig. 5g**). DDX55 was then overexpressed through transient transfection (**Fig. 5i and Supplementary Fig. 13b**), which significantly increased the activity of the reporter (**Fig. 5j**) in both cell lines, confirming that DDX55 overexpression leads to reduced miR-29b-3p activity. We did not observe these trends for miR-29a-3p and miR-29c-3p (**Supplementary Fig. 13c-f**). We further tested the impact of DDX55 knockdown on miR-29-3p function by measuring the protein level of Cyclin Dependent Kinase 6 (CDK6), a well-known target of miR-29-3p ^32–35^. CDK6 levels were reduced upon DDX55 suppression, which was rescued by inhibiting miR-29b-3p (**Fig. 5h**). DDX55 inhibition also substantially dampened cell viability (**Fig. 5k**), consistent with the previously mentioned difficulty in obtaining viable knockout clones. Taken together, these results confirm the inhibitory role of DDX55 on miR-29b-3p activity, a previously unrecognized mechanism that contributes to cancer progression.

## DISCUSSION

Leveraging the integration of mRNA stability profiles from 468 RBP inhibition experiments with predicted miRNA binding networks and following validation in a variety of external datasets, we identified new regulatory connections between specific RBPs and miRNAs that contribute to cancer transcriptome remodeling. Specifically, we show that well-established tumor suppressor miRNAs (e.g. miR-29-3p) and oncogenic miRNAs (e.g. miR-17-5p) are regulated by several RBPs across multiple cancer types, advancing our current knowledge about the mechanisms that underlie miRNA dysregulation in cancer.

The computational framework that we employed to infer changes in miRNA activity and RBP-miRNA connections presents noteworthy advantages. First, inferring miRNA activity from mRNA stability estimates derived from RNA-seq data avoids the need to perform small RNA sequencing (smRNA-seq), a specialized sequencing technique required to measure miRNA abundance on a large scale. As such, this approach enables broader use of the wealth of publicly available RNA-seq datasets, at virtually no additional costs. Second, while databases such as TCGA do provide miRNA expression data, abundance may not reflect the actual activity of miRNAs, which can be affected, for example, by the expression of targets, accessibility of binding sites, and interactions with other ncRNAs and RBPs. Previous evidence has indeed confirmed that the expression of miRNAs does not necessarily correlate with their activity ^36^. Our results extend on these previous reports by showing that changes in miRNA abundance cannot explain most of the variations observed in miRNA activity profiles in TCGA data. Third, our approach is designed to identify changes in activity of miRNAs based on dynamics in the stability of a large number of their target mRNAs. Thus, our method differs from those applied in previous reports that inferred miRNA activity from the coordinated changes in the abundance of their targets ^36^. This is important, since miRNAs act mostly at the post-transcriptional level. Therefore, inferring miRNA activity based on stability of target mRNAs, rather than their abundance, benefits from avoiding the confounding impacts of transcriptional changes. These benefits helped us replicate previous findings about the function of specific miRNAs in different cancers, including those of miR-30-5p in KIRC ^22–24^, miR-122-5p in CHOL and LIHC ^26,27^, and the pan-cancer dysregulation of miR-29-3p ^28^, miR-19-3p and miR-17-5p ^29,30^. Fourth, our approach is not restricted to a specific mechanism of regulation of miRNAs by RBPs, such as biogenesis or processing, or direct binding to mRNA targets. Accordingly, our method can detect any functional RBP-miRNA connections, whether direct or indirect, occurring at any step in the miRNA life cycle, from transcription to mRNA target repression. This feature distinguishes our approach from previous reports that focused on the impact of direct interactions between RBPs and miRNAs on the biogenesis of miRNAs, their binding to targets, (through *in vitro* pull-down assays) ^13^, eCLIP experiments ^14^, or transcriptome-wide screens of RBP and miRNA binding sites ^15^. We believe that our ‘process-agnostic’ approach is not only able to identify funtional RBP-miRNA connections that are linked to known biological processes, including miRNA biogenesis and miRNA binding to targets (e.g. PTBP1-miR-29) ^13,37^, but also captures funtional regulatory RBP-miRNA interactions without the need for prior knowledge about the nature of the interations. This computational framework enables the identification of new regulatory processes that are not easily identifiable by mechanism-specific approaches, including our discovery of the DDX55-miR-29-3p axis.

It should be noted that the predicted miRNA regulatory network used here has some limitations. First, only a limited number of human miRNAs were included (153 conserved miRNA families), representing only a subset of the thousands of miRNAs identified in humans to date. miRNA-mRNA interactions were also predicted based on seed matches to the 3’UTR; as such, non-canonical binding and miRNA binding to the 5’UTR or coding regions may lead to false-negatives in the network ^3,38^. Furthermore, these predictions cannot take the accessibility of binding sites into account, or the cell type specificity of the interactions. These limitations may lead to a reduced sensitivity for the inference of miRNA activity changes. Nevertheless, our study presents many novel functional regulatory connections between RBPs and miRNAs, highlighting RBPs that regulate the function of miRNAs with important roles in cancer.

Of particular interest, the inhibitory interaction between DDX55 and miR-29-3p was among the most robust observations we identified, with evident pan-cancer relevance. The pertinence of this interaction in cancer was highlighted by the marked upregulation of DDX55 expression and a global downregulation of miR-29-3p activity across most cancer types, which were highly correlated across TCGA tumors. DDX55 is an ATP-dependent helicase which was shown to associate with pre-ribosomes ^39^. However, there appears to be insufficient information on this RBP in the literature. Although recent work has implicated DDX55 in hepatocellular carcinoma progression ^40^ and bioinformatic studies have highlighted associations with metastasis and poor prognosis in colorectal and lung cancer ^41,42^, its broader role in tumorigenesis, potential contributions across cancer types, and functions in RNA regulation remain poorly characterized. In contrast, miR-29-3p has been extensively studied in the context of cancer. Several miR-29-3p targets are oncogenes or anti-apoptotic genes. Previous studies have shown that miR-29-3p acts as tumor suppressor and is downregulated in the majority of human cancers, regulating key oncogenic processes including cell proliferation, differentiation, apoptosis, metastasis, angiogenesis, and several others ^28^. miR-29-3p downregulation is associated with more aggressive tumors, metastasis, invasion and poor prognosis, while pharmaceutical restoration of miR-29-3p expression may ameliorate patient outlook ^34^. Overall, our results show that DDX55 expression is increased in cancer, associates with poor prognosis, and represses miR-29-3p. Having validated the functional connection between DDX55 and miR-29-3p experimentally, together with the observation that DDX55 inhibition reduces the viability of cancer cell lines and the expression of the miR-29-3p target and cell cycle gene CDK6, our study implicates DDX55 as a potential oncogene and therapeutic target that may be modulated to restore the activity of miR-29-3p in cancer.

## METHODS

### Predicted miRNA regulons

The binding of miRNAs to their target mRNAs depends primarily on base-pairing of the “seed region”, a 6-8 nucleotides sequence at the 5’ end of the miRNA, with complementary sequences located mainly in the 3’ UTR ^43^. miRNA binding maps were constructed by counting the number of exact miRNA seed matches in mRNA 3’ UTRs. 3’ UTR sequences of protein-coding genes were identified using the Ensembl GRCh38 version 87 genome annotation and restricted to the first 2000bp, while miRNA seed sequences were retrieved from TargetScan 7.2^44^,. To avoid the possible confounding effects of alternative splicing in the 3’UTR, miRNA binding site predictions were limited to genes where a single 3’UTR, composed of a single exon, was shared across all isoforms. To reduce the number of miRNAs, we selected the 153 miRNA families that are conserved across vertebrates (conservation score ≥1), as these miRNAs are more likely to regulate core cellular functions (**Supplementary Table 2**).

### AGO2-CLASH

To confirm direct interactions between selected miRNAs and predicted targets *in vivo*, we used AGO2-CLASH qRT-PCR assays. We used a CLIP-validated AGO2 antibody (anti-pan Ago antibody clone 2A8 (Millipore MABE56, 0.5mg/mL)) and followed a preparation protocol similar to AGO2 HITS-CLIP (Moore et al. Nature Protocols 2014). Concisely, samples were CIP- and PNK-treated on-bead following AGO2 immunoprecipitation, and proximity ligation was performed using T4 RNA ligase 1. RNA was phenol-chloroform extracted following treatment of samples with Proteinase K. Custom RT primers were used to generate cDNA, and miRNA-target primer-pairs based on each of the candidate interactions were used to perform qPCR (SYBR Green). Specifically, qPCR primers were designed using OligoCalc and Primer-BLAST, to amplify the hybrid formed by the ligation of a miRNA and its target. Briefly, the forward primer was designed to anneal to the miRNA part of the hybrid, while the reverse primer was designed to bind to the target transcript. A reverse transcription primer was also designed, to bind to the target transcript. Primer sequences can be found in **Supplementary Fig. 1a**. Negative control primers were also included, designed to capture the ligation of miRNAs and mRNAs which are not predicted to interact. Junctions that successfully amplified were marked as validated (**Supplementary Fig. 1b**).

### RNA-Seq datasets and processing

Raw RNA-seq data from shRNA-mediated knockdown of 230 different RBPs in HepG2 and/or K562 cell lines were obtained from ENCODE ^19^ (**Supplementary Table 1**). Raw data and gene expression data (Fragments per kilobase of transcript per million mapped reads; FPKM) from total RNA-sequencing of 308 cell line and tissue samples were also obtained from ENCODE (**Supplementary Table 1**). RNA-seq reads were mapped to the GRCh38 genome assembly using HISAT2 2.1.0^45^. We retrieved BAM files for 6903 tumor samples and 677 adjacent normal samples from the 17 cancer types with at least 5 normal samples in TCGA from the National Cancer Institute (NCI) Genomic Data Commons (GDC) data portal (https://portal.gdc.cancer.gov/GDC; dbGaP study accession phs000178.v1.p1). Glioblastoma (GBM) was excluded since the samples labelled as “normal” in this dataset do not represent normal adjacent tissue samples, but rather blood samples, contrary to the other cancer types. For the estimation of mRNA stability in all datasets, reads originating from exonic and intronic regions from the Ensembl GRCh38 version 87 GTF annotation and supported by both Ensembl and Havana consortia were quantified, as described previously ^31^. Briefly, to avoid the confounding effect of alternative splicing, we filtered for exonic reads mapping to constitutive exons. Intronic regions that overlapped an exon were excluded from our annotations, since it would be difficult to determine whether reads originating from these regions should be attributed to the exon or intron. RSeQC 2.6.4 ^46^ was then used to determine the strandedness of RNA-seq data and BAM files were sorted by read name using SAMtools 1.6. Exonic and intronic reads were counted separately using HTSeq-count 0.11.3^47^, using the “intersection-strict” mode for exonic reads and the “union” mode for intronic reads, and a MAPQ score cutoff ≥30. In the case of gene-level read counts, HTSeq-count was used with the “intersection-strict” mode based on original Ensembl GRCh38 v87 gene annotations. Gene-level normalized RNA-seq (RNA-Seq by Expectation-Maximization; RSEM) data (then converted to the log2 scale adding a pseudocount of 0.0001), mature miRNA abundance log2 reads per million mapped reads (RPM), and clinical data for TCGA datasets were obtained from Firebrowse (http://firebrowse.org/). For traditional differential gene expression analyses, DESeq2 1.46.0 ^48^ was used to compute differential gene expression.

### Inferring changes in mRNA stability and correcting for batch and bias

BETAS (https://github.com/csglab/BETAS) was used to estimate the changes in the abundances of pre-mRNA and mature mRNA for each gene, based on the changes in intronic and exonic read counts relative to the average of all samples, in the natural logarithmic scale (Δ*e* and Δ*i*) ^49^. BETAS estimates the distribution of abundance of exonic and intronic fragments of each gene from the observed exonic and intronic read counts using Bayesian statistics. More specifically, it uses Markov Chain Monte Carlo simulations with the Metropolis-Hastings algorithm to estimate the parameters of distributions, including their mean and variance. BETAS was also used to infer differential mRNA stability as the difference in the fold-change (also in the natural logarithm scale) of abundance of mature and pre-mRNA (Δ*e* − Δ*i*) ^49^. To prevent confounding from alternative splicing, we restricted the analysis to constitutive exons. Stability estimates were filtered by removing uncertain Δ*e* − Δ*i* estimates, i.e. estimates with high inferential variance as reported by BETAS. For the filtering, we selected the smallest standard deviation cut-off that retained at least 10,000 genes following the filtering procedure described below. For ENCODE datasets (RBP knockdown RNA-seq, cell line and tissue total RNA-seq), we first discarded genes with fewer than 20 values after filtering and then processed samples from each cell line separately. For TCGA datasets, samples with more than 20% missing values after filtering were discarded, as were genes for which more than 20% of normal or tumor samples had missing values.

Stability estimates for RBP knockdown experiments from ENCODE were batch-corrected, to account for technical artifacts from sample collection and processing. To define batches, correlation matrices were generated while setting Pearson correlation coefficients < 0 to 0, to only retain the samples that were positively correlated with each other (**Supplementary Fig. 3**). Hierarchical clustering using average linkage produced relatively homogeneous clusters around the diagonal. The dendrogram was manually analyzed to identify broad clusters that mostly had the same release date (and sometimes the same associated control, since these are related to the experiment batch) but consisted of various RBPs, and such clusters were deemed to represent experiments with similar batch effects. At least 20 experiments were included in most batches (**Supplementary Table 1**).

We further improved mRNA stability estimates using recent approaches that explicitly model the kinetics of mRNA metabolism, providing a more accurate estimate of changes in mRNA stability. A bias introduced by the maximum capacity of the pre-mRNA processing machinery and changes in transcription rate was accounted for by modeling the raw differential stability estimates as a function of intronic changes across samples ^16^, for each gene individually:

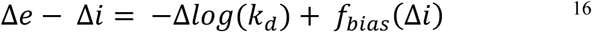

while also considering the batch in the case of the RBP knockdown experiments from ENCODE:

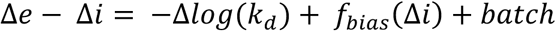

Here, *k*_d_ represents the mRNA decay constant, which represents the inverse of mRNA stability, and *f*_bias_ is the gene-specific bias function, which depends on the rate of transcription. Following Alkallas et al. (2017), we approximated *f*_bias_(Δ*i*) as a linear function, i.e. *f*_bias_(Δ*i*) = *β*_bias_ × Δ*i*. We corrected for this bias, together with the batch effects, by including the batch identification and intronic changes (Δ*i*) across samples as covariates in a robust linear regression (Δ*e* − Δ*i* ∼ *β*_bias_ × Δ*i* + *β*_batch(_ × *batc*ℎ + *β*_0_ + *ε*) and retaining the residuals, which were considered as an estimate of bias-and batch-corrected differential mRNA stability −Δ*log*(*k*_d_) for each gene (**Supplementary Fig. 2a and 3**). Robust linear regression, performed using the *rlm* function of the MASS R package (version 7.3. 60.2), was used due to its robustness to outliers and to violations of assumptions ^50^.

For TCGA datasets, since the transcription rate may be correlated with the tumor status, we included the sample type *sType* (tumor or normal) as a covariate in a similar linear model (Δ*e* − Δ*i* ∼ *β*_bias_ × Δ*i* + *β*_sType-_ × *sType* + *β*_0_ + *ε*). We then computed the residuals using a robust linear regression as described above, and added back the effect of the *sType* variable, to obtain partial residuals, which represent differential stability estimates in each tumor relative to the average of normal samples (**Supplementary Fig. 2b**).

### miRNA activity inference

To infer changes in miRNA activity, we used an approach derived from recently published methods ^16,17^. The dependence of differential mRNA stability on the presence of predicted miRNA binding sites in the 3’UTR was evaluated using a multivariate generalized linear model (‘glm’ function in R). mRNA properties that can affect stability, such as 3’UTR length, nucleotide and dinucleotide composition ^51^, were included as confounding variables in the model. Regression coefficients for each miRNA therefore represent how its binding to mRNAs can predict their differential stability. In other words, the coefficients represent the differential stability of miRNA targets. Since miRNA activity is inversely correlated with target differential stability, the coefficient signs were reversed to obtain miRNA activity profiles. miRNAs whose inferred binding was a significant predictor of differential mRNA stability were identified at false discovery rate (FDR < 0.05), using the Benjamini-Hochberg procedure.

### Correcting mRNA stability estimates for shRNA off-target effects

From the systematic RBP perturbation data from ENCODE, we had originally identified 59 inferred interactions between specific RBPs and miRNAs, which were statistically significant in at least one biological replicate at FDR < 0.05, with other replicates in at least one cell line being consistent in terms of direction of the effect (**Supplementary Fig. 4**). Most RBPs implicated appeared to act as negative miRNA regulators, with an increased activity of specific miRNAs upon RBP knockdown. This pattern, in addition to the observation that several of the shRNAs used to perform the knockdown experiments contained the seed sequence of the specific miRNA that were inferred to interact with the perturbed RBP (**Supplementary Fig. 5a**), led us to suspect that shRNA-off target effects might mimic miRNA function and lead to false positive changes in inferred miRNA activity. In other words, an shRNA that mediates off-target effects and mimics a miRNA in a particular knockdown sample would be detected by our method as an apparent increase in miRNA activity (upon RBP inhibition), and we would therefore infer an inhibitory role for the RBP on the miRNA. Off-target transcript silencing is a well-known unintended consequence of shRNA-mediated knockdown, is sequence dependent and arises from partial complementarily of the sense or antisense strands of the shRNA to an unexpected target. Furthermore, off-target silencing was shown to occur in a manner analogous to miRNA-mediated silencing ^52,53^. In such cases, regulatory effects of RBP on miRNA activity is indistinguishable from off-target effects of the shRNA.

To correct our inferences for potential off-target effects, we first identified the shRNA nucleotide positions most strongly associated with changes in mRNA stability across shRNAs (**Supplementary Fig. 5b**), which should represent the positions that mediate off-target silencing and mimic miRNA function. To do so, we tested the association between counts of each shRNA 6-mers or 7-mers generated using a sliding window in each gene’s 3’UTR and their change in mRNA stability, using a linear model. FDR correction was applied and shRNA positions that were significantly associated with off-target effects in more cases (samples) than expected by chance were identified using Fisher’s exact test. Stability measurements were then corrected for off-target effects from these specific positions. More specifically, 6-mer and 7-mer counts from these positions were included as confounders in a linear model testing their association with mRNA stability profiles, and residuals were retained as corrected stability data. Corrected mRNA stability profiles were then used for a second inference of differential miRNA activity as described above. To generate a new list of candidate RBP-miRNA interactions, all original candidate interactions were considered, and the new P-values of differential miRNA activity estimates of all corresponding samples were corrected for FDR and filtered at FDR < 0.05. New inferred candidate interactions with consistent interaction type (activating/inhibitory) across replicates and the original inference were retained as candidate interactions. We retained 39 candidate interactions, which are shown in **Fig. 2 a-b and Supplementary Table 3**.

### Generation of CRISPR-Cas9 knockout cell lines

HEK293T cells were cultured in Dulbecco’s modified Eagle medium (DMEM), along with 10% of fetal bovine serum (FBS) supplemented with HEPES and gentamycin. sgRNAs (**Supplementary Table 4**) were designed using ChopChop (https://chopchop.cbu.uib.no/) and inserted into the GFP-containing plasmid PX458 (Addgene plasmid # 48138) using BbsI restriction sites. Lipofectamine 2000 (Invitrogen) was used to transfect 2 ug of plasmid into the HEK293T cells in a 6-well plate, following the manufacturer’s instructions. GFP expression was used 2-3 days after transfection to sort cells, as single cells, into a 96-well plate. Following colony expansion, candidates were screening for knockout by Western blotting with antibodies against DDX55 (Invitrogen PA5-60612), PTBP1 (Invitrogen 32-5000) and SUPV3L1 (Bethyl Laboratories A303-056A). CRISPR targeted regions were amplified with PCR from the Genomic DNA of candidate clones, and sub-cloned using StrataClone (Agilent) for Sanger sequencing with the T7 primer (5’-TAATACGACTCACTATAGGG-3’) to confirm mutations. See **Supplementary Fig. 6** for clone selection.

### Inference of differential miRNA transcription in TCGA tumors

To investigate the relationship between transcription of miRNA host genes and the abundance of mature miRNAs, we interrogated miRNA loci included inside introns. For such miRNA genes, transcriptional changes can be approximated from that of the host gene, which can be estimated from changes in intronic read counts ^49^. Pri-miRNA genes annotation was obtained from miRbase (ftp://mirbase.smith.man.ac.uk/pub/mirbase/CURRENT/genomes/). Introns were defined from the Ensembl GRCh38 v87 annotation and those supported by both Ensembl and Havana consortia were included. Intronic regions overlapping any exon were excluded from the annotations. Pri-miRNA genes fully contained within these intronic regions were retained. We then retained host genes for which changes in abundance relative to the average were due at least in part to changes in transcription, as genes where changes in intronic read counts were significantly positively correlated (at FDR < 0.05) with changes in exonic read counts across at least 80% of TCGA cancers (**Supplementary Fig. 9a**). We then used changes in host gene transcription as a proxy for changes in miRNA transcription.

### Correcting for tumor purity

For the TCGA data, correction for sample purity was performed where applicable by including, as a confounding variable in the models, the consensus measurements of purity estimations (CPEs) ^54^, which represent combined measures of purity derived from two widely used algorithms, ESTIMATE and LUMP ^54,55^.

### Cell culture for DDX55 and miR-29-3p inhibition and overexpression experiments

The liver cancer cell line HepG2 and renal cancer cell line 786-O were obtained from the American Type Culture Collection (ATCC; Rockville, MD, USA) and cultured at 37^°^C with 5% CO_2_ in DMEM, following the provider’s protocol.

### DDX55 and miR-29-3p overexpression and inhibition studies

miRNA inhibition and overexpression using mimics, in addition to DDX55 siRNA-mediated knockdown, were performed in 2 X 10^5^ 786-O cells and 3 X 10^5^ HepG2 cells per well in a 6-well plate. Either 10 nM siGENOME Human DDX55 siRNA (Dharmacon #D-027082-03 and D-027082-17), AllStar negative control siRNA (Qiagen #1027281), hsa-miR-29b-3p *mir*Vana miRNA mimic (Ambion #4464066), hsa-miR-29a-3p *mir*Vana miRNA mimic (Life Technologies #4464066-MC12499), hsa-miR-29c-3p *mir*Vana miRNA mimic (Life Technologies #4464066-MC10518), *mir*Vana miRNA mimic negative control (Ambion #4464058), *mir*Vana miRNA inhibitor negative control (Ambion #4464076), hsa-miR-29b-3p *mir*Vana miRNA inhibitor (Ambion #4464084) were reverse-transfected using Lipofectamine RNAiMAX (Invitrogen) in antibiotic-free media, following the manufacturer’s protocol. After 8h, the media was replaced with growth media containing antibiotics, followed by incubation of the cells in a humidified incubator with 5% CO_2_ for 48 hrs or 72 hrs, at 37^°^C.

DDX55 overexpression was performed using 500 ng of DDX55 expression plasmid (OriGene C121509) or *pCMV6-XL5* empty vector (OriGene #PCMV6XL5) and reverse transfected with Lipofectamine 3000 (Invitrogen) as per the provider’s instructions. Transfected cells were incubated in a humidified incubator with 5% CO_2_ for 48 hrs or 72 hrs, at 37^°^C.

### Library preparation and RNA-sequencing

For selected CRISPR-mediated RBP KO clones, total RNA was extracted using QIAzol with the miRNeasy kit (Qiagen) following the provider’s recommendations, and sequencing was performed using Illumina NovaSeq Sprime (100bp paired-end).

### Western Blot Analysis

For the CRISPR-mediated RBP KOs, cells were lysed using lysis buffer (50 mM Tris-HCl pH 7.4, 150 mM NaCl, 1% sodium deoxycholate, 1% IGEPAL CA-630, 0.1% sodium dodecyl sulphate, and 1 mM ethylenediaminetetraacetic acid) with the addition of protease and phosphatase inhibitors (Sigma and Roche). Proteins from total cell lysates were immobilized on PVDF membranes (Millipore) and incubated first with DDX55, PTBP1, and SUPV3L1 antibodies at a 1:1,000 dilution in Odyssey blocking buffer (Li-COR), and then with anti-mouse or anti-rabbit secondary IR dye antibodies (Li-COR) at 1:10,000. The Li-COR imaging system was used to scan blots.

For DDX55 and miR-29 overexpression and inhibition experiments, protein extraction and western blotting for cell lines were performed as previously described ^17^. Signals were detected using Pierce^™^ ECL 2 Western Blotting Substrate (Thermo Fisher Scientific). The signal acquisition from the membranes and image analysis of the blots were carried out by exposing the membrane on ChemiDoc^™^ Imaging System powered by Image Lab^™^ software (Bio-Rad). The primary antibodies were as follows: DDX55 (Thermo Fisher Scientific #PA5-60612), CDK6 (Cell Signaling #3136) and GAPDH (Thermo Fisher Scientific #MA5-15738-HRP).

### Dual Luciferase Reporter Assays

Synthetic renilla luciferase *miRNA Target GoClone* vectors and *LightSwitch^™^ EMPTY-3′UTR* from SwitchGear Genomics were used as reporter vectors. Firefly luciferase, *pCI-neo-firefly-luc* (Promega), was used as the internal control vector. The effect of RBPs on the activity of miRNAs was investigated using Renilla-reporter and firefly-internal control dual luciferase reporter assays. A total of 3 X 10^4^ 786-O cells and 4 X 10^4^ HepG2 cells were placed in in antibiotic-free media, in each well of a 24-well plate and reverse transfected with 10 nM of siRNAs or miRNA mimics, and 100 ng of DDX55 expressing or control plasmids. The media was replaced with media containing antibiotics after 8 hrs and cells were incubated in a humidified incubator with 5% CO_2_ for 24 hrs at 37^°^C. Growth media was removed the following day and cells were washed with PBS for forward-transfection with 20 ng of renilla vectors and 4 ng of firefly vectors using Lipofectamine 3000 (Invitrogen), following to the manufacturer’s recommendations. The Dual-Luciferase^®^ Reporter Assay System (Promega) was used to test for luciferase activity, two days after transfection with reporter vectors and following the provider’s protocol. Cells were lysed and transferred to Nunc^™^ F96 MicroWell^™^ White Polystyrene luminometer plates (Thermo Fisher Scientific). TECAN Infinite^®^ M200 PRO microplate reader with Te-Inject^™^ reagent injectors (Tecan Group, Ltd., Switzerland) were used for luciferase measurements, carried out in five replicates in three independent experiments. Renilla luciferase measurements were first normalized to the firefly luciferase activity, and the relative luciferase activity was presented.

### Cell viability

Cell viability was assessed using a sulforhodamine B sodium salt (SRB) (Sigma-Aldrich) assay, following the manufacturer’s instructions.

### Statistics and reproducibility

All statistical analyses were performed using Bioconductor packages in R (version 4.4.1). The specific statistical tests used for each analysis and the associated measures of statistical significance are indicated within the main text, methods, in the figure, or in their legends. Statistical significance was set at P < 0.05 for all analyses and multiple testing correction was performed when applicable. Adjusted P-values were calculated using the Benjamini-Hochberg false discovery rate procedure. Sample size for TCGA cohort analysis depended on publicly available data. To ensure reproducibility for RNA-seq experiments, biological replicates were used and/or the findings were replicated in other cell lines.

## DATA AVAILABILITY

All data generated during this study are included in this published article (and its supplementary information files). Supplementary Data files and additional analysis files are available via Zenodo at https://doi.org/10.5281/zenodo.22679902. RNA-seq data from the RBP CRISPR-mediated knockout experiments is available via GEO under accession GSE202459. Other data used in this paper are available via their source publications as indicated in the article. The results shown here are in part based upon data generated by the TCGA Research Network: https://www.cancer.gov/tcga.

## CODE AVAILABILITY

BETAS is available at https://github.com/csglab/BETAS. The code used to perform the analyses and generate the figures is staged on GitHub at https://github.com/csglab/RBP-miRNA-manuscript.

## ACKNOWLEDGEMENTS

This work was supported by McGill University and funds from the Canadian Institutes of Health Research (CIHR; PJT-155966) to H.S.N.. The work is also supported by resource allocations from Compute Canada to H.S.N. H.S.N holds a Canada Research Chair funded by the CIHR. G.P. and R.A. hold training scholarships from the CIHR, Oncopole and the Fonds de recherche du Québec – Santé (FRQS). Y.R. is supported by the FRQS. The results published here are in part based on data generated by the TCGA Research Network: https://www.cancer.gov/tcga.

## AUTHOR CONTRIBUTIONS

G.P., H.S.N. and Y.R. conceived the study. G.P. and H.S.N. developed the computational methods. G.P. analyzed the data and wrote the manuscript, with edits from H.S.N and Y.R.. M.A., C.T., P.J., E.M., T.N., M.R., B.C. and K.G. performed the experiments. R.A. contributed to data processing. H.G. and T.D. contributed to experimental design. I.R.W contributed to manuscript revision. Y.R. and H.S.N. supervised all research.

## Footnotes

1 We use miR-17-5p to refer to the miR-17-5p/20-5p/93-5p/106-5p/519-3p family.

## REFERENCES

1 Garraway, L. A. & Lander, E. S. Lessons from the cancer genome. Cell 153, 17–37, doi:10.1016/j.cell.2013.03.002 (2013).

2 Vogelstein, B. et al. Cancer genome landscapes. Science 339, 1546–1558, doi:10.1126/science.1235122 (2013).

3 Friedman, R. C., Farh, K. K., Burge, C. B. & Bartel, D. P. Most mammalian mRNAs are conserved targets of microRNAs. Genome Res 19, 92–105, doi:10.1101/gr.082701.108 (2009).

4 Jansson, M. D. & Lund, A. H. MicroRNA and cancer. Mol Oncol 6, 590–610, doi:10.1016/j.molonc.2012.09.006 (2012).

5 Hayes, J., Peruzzi, P. P. & Lawler, S. MicroRNAs in cancer: biomarkers, functions and therapy. Trends Mol Med 20, 460–469, doi:10.1016/j.molmed.2014.06.005 (2014).

6 Gerstberger, S., Hafner, M. & Tuschl, T. A census of human RNA-binding proteins. Nat Rev Genet 15, 829–845, doi:10.1038/nrg3813 (2014).

7 Van Nostrand, E. L. et al. A large-scale binding and functional map of human RNA-binding proteins. Nature 583, 711–719, doi:10.1038/s41586-020-2077-3 (2020).

8 Loffreda, A., Rigamonti, A., Barabino, S. M. & Lenzken, S. C. RNA-Binding Proteins in the Regulation of miRNA Activity: A Focus on Neuronal Functions. Biomolecules 5, 2363–2387, doi:10.3390/biom5042363 (2015).

9 Iadevaia, V. & Gerber, A. P. Combinatorial Control of mRNA Fates by RNA-Binding Proteins and Non-Coding RNAs. Biomolecules 5, 2207–2222, doi:10.3390/biom5042207 (2015).

10 Wang, Z. L. et al. Comprehensive Genomic Characterization of RNA-Binding Proteins across Human Cancers. Cell Rep 22, 286–298, doi:10.1016/j.celrep.2017.12.035 (2018).

11 Kechavarzi, B. & Janga, S. C. Dissecting the expression landscape of RNA-binding proteins in human cancers. Genome Biol 15, R14, doi:10.1186/gb-2014-15-1-r14 (2014).

12 Han, J. et al. Multilayered control of splicing regulatory networks by DAP3 leads to widespread alternative splicing changes in cancer. Nat Commun 13, 1793, doi:10.1038/s41467-022-29400-7 (2022).

13 Treiber, T. et al. A Compendium of RNA-Binding Proteins that Regulate MicroRNA Biogenesis. Mol Cell 66, 270–284 e213, doi:10.1016/j.molcel.2017.03.014 (2017).

14 Nussbacher, J. K. & Yeo, G. W. Systematic Discovery of RNA Binding Proteins that Regulate MicroRNA Levels. Mol Cell 69, 1005–1016 e1007, doi:10.1016/j.molcel.2018.02.012 (2018).

15 Kim, S. et al. The regulatory impact of RNA-binding proteins on microRNA targeting. Nat Commun 12, 5057, doi:10.1038/s41467-021-25078-5 (2021).

16 Alkallas, R., Fish, L., Goodarzi, H. & Najafabadi, H. S. Inference of RNA decay rate from transcriptional profiling highlights the regulatory programs of Alzheimer’s disease. Nat Commun 8, 909, doi:10.1038/s41467-017-00867-z (2017).

17 Perron, G. et al. A General Framework for Interrogation of mRNA Stability Programs Identifies RNA-Binding Proteins that Govern Cancer Transcriptomes. Cell Rep 23, 1639–1650, doi:10.1016/j.celrep.2018.04.031 (2018).

18 Helwak, A. & Tollervey, D. Mapping the miRNA interactome by cross-linking ligation and sequencing of hybrids (CLASH). Nat Protoc 9, 711–728, doi:10.1038/nprot.2014.043 (2014).

19 Consortium, E. P. An integrated encyclopedia of DNA elements in the human genome. Nature 489, 57–74, doi:10.1038/nature11247 (2012).

20 Liu, Y. et al. miR-19a promotes colorectal cancer proliferation and migration by targeting TIA1. Mol Cancer 16, 53, doi:10.1186/s12943-017-0625-8 (2017).

21 Jiang, T. et al. miR-19b-3p promotes colon cancer proliferation and oxaliplatin-based chemoresistance by targeting SMAD4: validation by bioinformatics and experimental analyses. J Exp Clin Cancer Res 36, 131, doi:10.1186/s13046-017-0602-5 (2017).

22 Chen, Z. et al. The putative tumor suppressor microRNA-30a-5p modulates clear cell renal cell carcinoma aggressiveness through repression of ZEB2. Cell Death Dis 8, e2859, doi:10.1038/cddis.2017.252 (2017).

23 Huang, J. et al. Hypoxia-induced downregulation of miR-30c promotes epithelial-mesenchymal transition in human renal cell carcinoma. Cancer Sci 104, 1609–1617, doi:10.1111/cas.12291 (2013).

24 Song, S. et al. Urinary exosome miR-30c-5p as a biomarker of clear cell renal cell carcinoma that inhibits progression by targeting HSPA5. J Cell Mol Med 23, 6755–6765, doi:10.1111/jcmm.14553 (2019).

25 Jopling, C. Liver-specific microRNA-122: Biogenesis and function. RNA Biol 9, 137–142, doi:10.4161/rna.18827 (2012).

26 Liu, N. et al. The Roles of MicroRNA-122 Overexpression in Inhibiting Proliferation and Invasion and Stimulating Apoptosis of Human Cholangiocarcinoma Cells. Sci Rep 5, 16566, doi:10.1038/srep16566 (2015).

27 Nakao, K., Miyaaki, H. & Ichikawa, T. Antitumor function of microRNA-122 against hepatocellular carcinoma. J Gastroenterol 49, 589–593, doi:10.1007/s00535-014-0932-4 (2014).

28 Kwon, J. J., Factora, T. D., Dey, S. & Kota, J. A Systematic Review of miR-29 in Cancer. Mol Ther Oncolytics 12, 173–194, doi:10.1016/j.omto.2018.12.011 (2019).

29 Olive, V., Li, Q. & He, L. mir-17-92: a polycistronic oncomir with pleiotropic functions. Immunol Rev 253, 158–166, doi:10.1111/imr.12054 (2013).

30 He, L. et al. A microRNA polycistron as a potential human oncogene. Nature 435, 828–833, doi:10.1038/nature03552 (2005).

31 Perron, G. et al. Pan-cancer analysis of mRNA stability for decoding tumour post-transcriptional programs. Commun Biol 5, 851, doi:10.1038/s42003-022-03796-w (2022).

32 Zhao, J. J. et al. microRNA expression profile and identification of miR-29 as a prognostic marker and pathogenetic factor by targeting CDK6 in mantle cell lymphoma. Blood 115, 2630–2639, doi:10.1182/blood-2009-09-243147 (2010).

33 Zhu, K. et al. MiR-29b suppresses the proliferation and migration of osteosarcoma cells by targeting CDK6. Protein Cell 7, 434–444, doi:10.1007/s13238-016-0277-2 (2016).

34 Garzon, R. et al. MicroRNA 29b functions in acute myeloid leukemia. Blood 114, 5331–5341, doi:10.1182/blood-2009-03-211938 (2009).

35 Li, Y. et al. Progressive miRNA expression profiles in cervical carcinogenesis and identification of HPV-related target genes for miR-29. J Pathol 224, 484–495, doi:10.1002/path.2873 (2011).

36 Lee, E. et al. Inferred miRNA activity identifies miRNA-mediated regulatory networks underlying multiple cancers. Bioinformatics 32, 96–105, doi:10.1093/bioinformatics/btv531 (2016).

37 Xue, Y. et al. Direct conversion of fibroblasts to neurons by reprogramming PTB-regulated microRNA circuits. Cell 152, 82–96, doi:10.1016/j.cell.2012.11.045 (2013).

38 Martin, H. C. et al. Imperfect centered miRNA binding sites are common and can mediate repression of target mRNAs. Genome Biol 15, R51, doi:10.1186/gb-2014-15-3-r51 (2014).

39 Choudhury, P., Kretschmer, J., Hackert, P., Bohnsack, K. E. & Bohnsack, M. T. The DExD box ATPase DDX55 is recruited to domain IV of the 28S ribosomal RNA by its C-terminal region. RNA Biol 18, 1124–1135, doi:10.1080/15476286.2020.1829366 (2021).

40 Yu, B. et al. DDX55 promotes hepatocellular carcinoma progression by interacting with BRD4 and participating in exosome-mediated cell-cell communication. Cancer Sci 113, 3002–3017, doi:10.1111/cas.15393 (2022).

41 Cui, Y. et al. Lead DEAD/H box helicase biomarkers with the therapeutic potential identified by integrated bioinformatic approaches in lung cancer. Comput Struct Biotechnol J 19, 261–278, doi:10.1016/j.csbj.2020.12.007 (2021).

42 Liu, J. et al. Identification of liver metastasis-associated genes in human colon carcinoma by mRNA profiling. Chin J Cancer Res 30, 633–646, doi:10.21147/j.issn.1000-9604.2018.06.08 (2018).

43 Lewis, B. P., Shih, I. h., Jones-Rhoades, M. W., Bartel, D. P. & Burge, C. B. Prediction of Mammalian MicroRNA Targets. Cell 115, 787–798, doi:10.1016/s0092-8674(03)01018-3 (2003).

44 Agarwal, V., Bell, G. W., Nam, J. W. & Bartel, D. P. Predicting effective microRNA target sites in mammalian mRNAs. Elife 4, doi:10.7554/eLife.05005 (2015).

45 Kim, D., Paggi, J. M., Park, C., Bennett, C. & Salzberg, S. L. Graph-based genome alignment and genotyping with HISAT2 and HISAT-genotype. Nat Biotechnol 37, 907–915, doi:10.1038/s41587-019-0201-4 (2019).

46 Wang, L., Wang, S. & Li, W. RSeQC: quality control of RNA-seq experiments. Bioinformatics 28, 2184–2185, doi:10.1093/bioinformatics/bts356 (2012).

47 Anders, S., Pyl, P. T. & Huber, W. HTSeq--a Python framework to work with high-throughput sequencing data. Bioinformatics 31, 166–169, doi:10.1093/bioinformatics/btu638 (2015).

48 Love, M. I., Huber, W. & Anders, S. Moderated estimation of fold change and dispersion for RNA-seq data with DESeq2. Genome Biol 15, 550, doi:10.1186/s13059-014-0550-8 (2014).

49 Gaidatzis, D., Burger, L. & Stadler, M. B. Analysis of intronic and exonic reads in RNA-seq data characterizes transcriptional and post-transcriptional regulation. Nat Biotechnol 33, 722–729, doi:10.1038/nbt.3269 (2015).

50 Huber, P. J. International Encyclopedia of Statistical Science. *Springer, Berlin*, Heidelberg, doi:10.1007/978-3-642-04898-2_594 (2011).

51 Wu, X. & Brewer, G. The regulation of mRNA stability in mammalian cells: 2.0. Gene 500, 10–21, doi:10.1016/j.gene.2012.03.021 (2012).

52 Jackson, A. L. et al. Widespread siRNA “off-target” transcript silencing mediated by seed region sequence complementarity. RNA 12, 1179–1187, doi:10.1261/rna.25706 (2006).

53 Lin, X. et al. siRNA-mediated off-target gene silencing triggered by a 7 nt complementation. Nucleic Acids Res 33, 4527–4535, doi:10.1093/nar/gki762 (2005).

54 Aran, D., Sirota, M. & Butte, A. J. Systematic pan-cancer analysis of tumour purity. Nat Commun 6, 8971, doi:10.1038/ncomms9971 (2015).

55 Yoshihara, K. et al. Inferring tumour purity and stromal and immune cell admixture from expression data. Nat Commun 4, 2612, doi:10.1038/ncomms3612 (2013).

56 Shannon, P. et al. Cytoscape: a software environment for integrated models of biomolecular interaction networks. Genome Res 13, 2498–2504, doi:10.1101/gr.1239303 (2003).

57 Korotkevich, G., et al. Fast gene set enrichment analysis. bioRxiv, doi:10.1101/060012 (2021).

58 McCarthy, D. J., Campbell, K. R., Lun, A. T. & Wills, Q. F. Scater: pre-processing, quality control, normalization and visualization of single-cell RNA-seq data in R. Bioinformatics 33, 1179–1186, doi:10.1093/bioinformatics/btw777 (2017).

