## Supplementary Information for "A map of RBP-miRNA regulatory connections for deciphering the mechanisms of transcriptome remodeling in cancer"

**Supplementary Table 1.** List of supplementary data

The Zenodo record is available at https://zenodo.org/records/22679902

Download URL: https://zenodo.org/records/22679902/files/code_input_data.zip?download=1

| |
|---|
| **File name:** Supplementary Data 1.xlsx<br>**Description: shRNA-mediated RBP knockdown experiments from ENCODE.** (**a**) Sample metadata and identified batches for shRNA-mediated RBP knockdown experiments from ENCODE. (**b**) Bias- and batch-corrected mRNA stability estimates for HepG2. Differential estimates are relative to the average of all samples and following correction for shRNA-off target effects. The column headers are in the following format: "<sample ID>_ <targeted RBP>_<biological replicate><technical replicate>_ <cell line>". (**c**) Same as (b), for K562. (**d**) Inferred differential miRNA activity. |
| **File name:** Supplementary Data 2.xlsx<br>**Description: ENCODE tissues and cell lines.** (**a**) Metadata for the total RNA-seq samples of tissues, primary and immortalized cell lines from ENCODE. (**b**) Bias-corrected mRNA stability estimates in ENCODE tissues and cell lines. Differential estimates are relative to the average of all samples. The column headers are in the following format: "<sample ID_<biological replicate>". (**c**) Inferred differential miRNA activity in ENCODE tissues and cell lines. Differential estimates are relative to the average of all samples. |
| **File name:** Supplementary Data 3.xlsx<br>**Description: Differential gene expression analysis in CRISPR-mediated RBP knockouts**. (a) DDX55, (b) PTBP1 and (c) SUPV3L1, relative to empty vector controls. |
| **File name:** Supplementary Data 4.xlsx<br>**Description: Bias-corrected mRNA stability estimates in TCGA tumors.** Differential estimates are relative to the average of normal samples. Individual tumor types are on separate sheets. |
| **File name:** Supplementary Data 5.xlsx<br>**Description: Inferred differential miRNA activity in TCGA tumors.** Estimates are relative to the average of normal samples. |
| **File name:** Supplementary Data 6.xlsx<br>**Description: Estimates of changes in mRNA abundance in the TCGA tumor samples.** Based on the changes in exonic read counts. Estimates are relative to the average of all samples. Individual tumor types are on separate sheets. |
| **File name:** Supplementary Data 7.xlsx<br>**Description: Estimates of changes in transcription in the TCGA tumor samples.** Based on the changes in intronic read counts. Estimates are relative to the average of all samples. Individual tumor types are on separate sheets. |

| miRNA family | Seed |
|---|---|
| let-7-5p/98-5p | GAGGUAG |
| miR-101-3p.1 | ACAGUAC |
| miR-103-3p/107 | GCAGCAU |
| miR-10-5p | ACCCUGU |
| miR-122-5p | GGAGUGU |
| miR-124-3p.1 | AAGGCAC |
| miR-125a-3p | CAGGUGA |
| miR-125-5p | CCCUGAG |
| miR-126-3p.1 | CGUACCG |
| miR-127-3p | CGGAUCC |
| miR-128-3p | CACAGUG |
| miR-129-5p | UUUUUGC |
| miR-130-3p/301-3p/454-3p | AGUGCAA |
| miR-132-3p/212-3p | AACAGUC |
| miR-133a-3p.2/133b | UUGGUCC |
| miR-134-5p | GUGACUG |
| miR-135-5p | AUGGCUU |
| miR-136-5p | CUCCAUU |
| miR-137 | UAUUGCU |
| miR-138-5p | GCUGGUG |
| miR-139-5p | CUACAGU |
| miR-140-5p | AGUGGUU |
| miR-141-3p/200a-3p | AACACUG |
| miR-142-3p.1 | GUAGUGU |
| miR-143-3p | GAGAUGA |
| miR-144-3p | ACAGUAU |
| miR-145-5p | UCCAGUU |
| miR-146-5p | GAGAACU |
| miR-148-3p/152-3p | CAGUGCA |
| miR-149-5p | CUGGCUC |
| miR-150-5p | CUCCCAA |
| miR-153-3p | UGCAUAG |
| miR-154-5p | AGGUUAU |
| miR-155-5p | UAAUGCU |
| miR-15-5p/16-5p/195-5p/424-5p/497-5p | AGCAGCA |
| miR-17-5p/20-5p/93-5p/106-5p/519-3p | AAAGUGC |
| miR-181-5p | ACAUUCA |
| miR-182-5p | UUGGCAA |
| miR-183-5p.1 | AUGGCAC |
| miR-184 | GGACGGA |
| miR-185-5p | GGAGAGA |
| miR-186-5p | AAAGAAU |
| miR-187-3p | CGUGUCU |
| miR-18-5p | AAGGUGC |
| miR-190-5p | GAUAUGU |
| miR-191-5p | AACGGAA |
| miR-192-5p/215-5p | UGACCUA |
| miR-193-3p | ACUGGCC |
| miR-194-5p | GUAACAG |
| miR-196-5p | AGGUAGU |
| miR-197-3p | UCACCAC |

| miRNA family | Seed |
|---|---|
| miR-199-5p | CCAGUGU |
| miR-19-3p | GUGCAAA |
| miR-1-3p/206 | GGAAUGU |
| miR-200bc-3p/429 | AAUACUG |
| miR-202-3p | GAGGUAU |
| miR-203a-3p.2 | UGAAAUG |
| miR-204-5p/211-5p | UCCCUUU |
| miR-205-5p | CCUUCAU |
| miR-208-3p | UAAGACG |
| miR-21-5p/590-5p | AGCUUAU |
| miR-210-3p | UGUGCGU |
| miR-214-3p/3619-5p | CAGCAGG |
| miR-216a-5p | AAUCUCA |
| miR-216b-5p | AAUCUCU |
| miR-217 | ACUGCAU |
| miR-218-5p | UGUGCUU |
| miR-219-5p | GAUUGUC |
| miR-22-3p | AGCUGCC |
| miR-221-3p/222-3p | GCUACAU |
| miR-223-3p | GUCAGUU |
| miR-224-5p | AAGUCAC |
| miR-23-3p | UCACAUU |
| miR-24-3p | GGCUCAG |
| miR-25-3p/32-5p/92-3p/363-3p/367-3p | AUUGCAC |
| miR-26-5p | UCAAGUA |
| miR-27-3p | UCACAGU |
| miR-28-5p/708-5p | AGGAGCU |
| miR-371-5p | CUCAAAC |
| miR-296-3p | AGGGUUG |
| miR-299-3p | AUGUGGG |
| miR-29-3p | AGCACCA |
| miR-381-3p | AUACAAG |
| miR-30-5p | GUAAACA |
| miR-31-5p | GGCAAGA |
| miR-320 | AAAGCUG |
| miR-324-5p | GCAUCCC |
| miR-326 | CUCUGGG |
| miR-328-3p | UGGCCCU |
| miR-329-3p/362-3p | ACACACC |
| miR-335-5p | CAAGAGC |
| miR-338-3p | CCAGCAU |
| miR-339-5p | CCCUGUC |
| miR-365-3p | AAUGCCC |
| miR-33-5p | UGCAUUG |
| miR-340-5p | UAUAAAG |
| miR-342-3p | CUCACAC |
| miR-346 | GUCUGCC |
| miR-34-5p/449-5p | GGCAGUG |
| miR-361-5p | UAUCAGA |
| miR-370-3p | CCUGCUG |
| miR-374-5p | UAUAAUA |

| miRNA family | Seed |
|---|---|
| miR-375 | UUGUUCG |
| miR-376-3p | UCAUAGA |
| miR-376c-3p | ACAUAGA |
| miR-377-3p | UCACACA |
| miR-378-3p | CUGGACU |
| miR-379-5p | GGUAGAC |
| miR-382-5p | AAGUUGU |
| miR-383-5p.1 | GAUCAGA |
| miR-384 | UUCCUAG |
| miR-410-3p | AUAUAAC |
| miR-411-5p.1 | AGUAGAC |
| miR-421 | UCAACAG |
| miR-425-5p | AUGACAC |
| miR-431-5p | GUCUUGC |
| miR-433-3p | UCAUGAU |
| miR-448 | UGCAUAU |
| miR-450-5p | UUUGCGA |
| miR-451 | AACCGUU |
| miR-455-5p | AUGUGCC |
| miR-485-5p | GAGGCUG |
| miR-486-5p | CCUGUAC |
| miR-487-3p | AUCGUAC |
| miR-488-3p | UGAAAGG |
| miR-490-3p | AACCUGG |
| miR-491-5p | GUGGGGA |
| miR-494-3p | GAAACAU |
| miR-495-3p | AACAAAC |
| miR-496.1 | GAGUAUU |
| miR-499a-5p | UAAGACU |
| miR-503-5p | AGCAGCG |
| miR-4725-5p | GACCCUG |
| miR-505-3p.1 | GUCAACA |
| miR-539-5p | GAGAAAU |
| miR-542-3p | GUGACAG |
| miR-543 | AACAUUC |
| miR-544a-3p | UUCUGCA |
| miR-551-3p | CGACCCA |
| miR-590-3p | AAUUUUA |
| miR-592/599 | UUGUGUC |
| miR-615-3p | CCGAGCC |
| miR-653-3p | UGUUGAA |
| miR-7-5p | GGAAGAC |
| miR-758-3p | UUGUGAC |
| miR-873-5p.1 | CAGGAAC |
| miR-874-3p | UGCCCUG |
| miR-875-5p | AUACCUC |
| miR-876-5p | GGAUUUC |
| miR-9-5p | CUUUGGU |
| miR-302-3p/372-3p/373-3p/520-3p | AAGUGCU |
| miR-96-5p/1271-5p | UUGGCAC |
| miR-99-5p/100-5p | ACCCGUA |

**Supplementary Table 2. Conserved miRNA families.** 153 miRNA families conserved across vertebrates, along with their seed sequences. See Methods for more detail.

| RBP | miRNA family | Type | Cell line |
|---|---|---|---|
| AATF | miR-455-5p | - | HepG2 |
| ABCF1 | miR-33a-3p | - | HepG2 |
| AKAP1 | miR-370-3p | - | K562 |
| BCCIP | miR-33-5p | - | HepG2 |
| DDX55 | miR-29-3p | - | HepG2 |
| DKC1 | miR-125a-3p | + | HepG2 |
| DKC1 | miR-592 | - | HepG2 |
| DNAJC2 | miR-21-5p/590-5p | - | HepG2 |
| EIF4B | miR-26-5p | - | HepG2 |
| FAM120A | miR-29-3p | + | HepG2 |
| FMR1 | miR-296-3p | + | HepG2 |
| FIP1L1 | miR-29-3p | - | HepG2 |
| IGF2BP1 | miR-758-3p | - | HepG2 |
| IGF2BP2 | miR-17-5p/20-5p/93-5p/106-5p/519-3p | + | HepG2 |
| ILF2 | miR-22-3p | + | HepG2 |
| MATR3 | miR-758-3p | - | HepG2 |
| MSI2 | miR-17-5p/20-5p/93-5p/106-5p/519-3p | + | HepG2 |
| MSI2 | miR-218-5p | - | HepG2 |
| NELFE | miR-96-5p/1271-5p | - | HepG2 |
| PABPC4 | miR-17-5p/20-5p/93-5p/106-5p/519-3p | + | HepG2 |
| PCBP1 | miR-27-3p | - | HepG2 |
| PSIP1 | miR-219-5p | - | HepG2 |
| PTBP1 | miR-29-3p | - | HepG2 |
| PUM1 | miR-25-3p/32-5p/92-3p/363-3p/367-3p | - | HepG2 |
| QKI | miR-124-3p.1 | - | HepG2 |
| QKI | miR-485-5p | - | HepG2 |
| RECQL | miR-96-5p/1271-5p | - | HepG2 |
| RPLP0 | miR-29-3p | - | HepG2 |
| RRP9 | miR-27-3p | - | HepG2 |
| RTF1 | miR-126-3p | - | HepG2 |
| SERBP1 | miR-133a-3p/133b | - | HepG2 |
| SF1 | miR-199-5p | - | HepG2 |
| STIP1 | miR-182-5p | - | HepG2 |
| SUCLG1 | miR-376-3p | - | HepG2 |
| SUGP2 | miR-23-3p | - | HepG2 |
| SUPV3L1 | miR-29-3p | + | HepG2 |
| SUPV3L1 | miR-17-5p/20-5p/93-5p/106-5p/519-3p | + | K562 |
| SUPV3L1 | miR-19-3p | + | K562 |
| YBX3 | miR-150-5p | + | HepG2 |

**Supplementary Table 3. Regulatory connections between RBPs and miRNAs, inferred from RBP inhibition experiments from ENCODE.** Regulatory pairs for which the change in miRNA activity upon RBP inhibition was significant (FDR < 0.05) and for which biological replicates were in agreement in terms of type of interaction (inhibitory or activating). The Type column represents the type of interaction, where a plus (+) sign represents an activating interaction and a minus (-) sign represents an inhibitory interaction. The cell line in which the interactions were inferred is also indicated.

| RBP | sgRNA | Sequence |
|---|---|---|
| PTBP1 | PTBP1-sg3-F | caccgGAAGCTCCCCATCGACGTCA |
| PTBP1 | PTBP1-sg3-R | aaacTGACGTCGATGGGGAGCTTCc |
| SUPV3L1 | SUPV3L1-sg1-F | caccgTGGAGCTTATGTACACAACG |
| SUPV3L1 | SUPV3L1-sg1-R | aaacCGTTGTGTACATAAGCTCCAc |
| DDX55 | DDX55-sg1-F | caccgCATGACGCCGGTGCAGGTAT |
| DDX55 | DDX55-sg1-R | aaacATACCTGCACCGGCGTCATGc |
| DDX55 | DDX55-sg3-F | caccgACTACTACATGGTAAGCGCC |
| DDX55 | DDX55-sg3-R | aaacGGCGCTTACCATGTAGTAGTc |

**Supplementary Table 4. Sequences of CRISPR sgRNAs.** List of sgRNAs for the CRISPR/Cas9-mediated knockdown of PTBP1, SUPV3L1 and DDX55.

| | |
|---|---|
| BLCA | let-7-5p/miR-98-5p |
| | miR-200bc-3p/429 |
| | miR-29-3p |
| BRCA | let-7-5p/miR-98-5p |
| | miR-143-3p |
| | miR-19-3p |
| | miR-200bc-3p/429 |
| | miR-21-5p/590-5p |
| | miR-26-5p |
| | miR-29-3p |
| CHOL | miR-122-5p |
| COAD | miR-138-5p |
| | miR-19-3p |
| | miR-25-3p/32-5p/92-3p/363-3p/367-3p |
| | miR-29-3p |
| | miR-590-3p |
| ESCA | miR-19-3p |
| | miR-29-3p |
| HNSC | miR-29-3p |
| KICH | miR-221-3p |
| KIRC | miR-142-3p.1 |
| | miR-29-3p |
| | miR-30-5p |
| | miR-590-3p |
| KIRP | miR-29-3p |
| | miR-30-5p |
| | miR-376c-3p |
| LIHC | miR-122-5p |
| LUAD | let-7-5p/miR-98-5p |
| | miR-138-5p |
| | miR-1-3p/206 |
| | miR-200bc-3p/429 |
| | miR-29-3p |
| | miR-30-5p |
| LUSC | let-7-5p/miR-98-5p |
| | miR-26-5p |
| | miR-29-3p |
| | miR-30-5p |
| PRAD | miR-29-3p |
| READ | miR-29-3p |
| STAD | let-7-5p/miR-98-5p |
| | miR-19-3p |
| | miR-200bc-3p/429 |
| | miR-29-3p |
| | miR-376c-3p |
| THCA | miR-29-3p |
| UCEC | let-7-5p/miR-98-5p |
| | miR-19-3p |
| | miR-200bc-3p/429 |
| | miR-204-5p/211-5p |
| | miR-26-5p |
| | miR-29-3p |
| | miR-9-5p |

**Supplementary Table 5. Differentially active miRNAs in TCGA tumors.** List of miRNA families with significant changes in inferred activity (FDR $< 0.05$) in at least 10 tumor samples in at least one TCGA cancer, relative to the average of normal samples. Red and blue indicate increased or reduced miRNA activity, respectively, based on the median change in activity.

| RBP | miRNA family | type |
|---|---|---|
| DDX55 | miR-29-3p | - |
| PTBP1 | miR-29-3p | - |
| RPLP0 | miR-29-3p | - |
| FAM120A | miR-29-3p | + |
| SF1 | miR-199-5p | - |
| PABPC4 | miR-17-5p/20-5p/93-5p/106-5p/519-3p | + |
| IGF2BP2 | miR-17-5p/20-5p/93-5p/106-5p/519-3p | + |
| MSI2 | miR-17-5p/20-5p/93-5p/106-5p/519-3p | + |
| SUPV3L1 | miR-17-5p/20-5p/93-5p/106-5p/519-3p | + |
| SUPV3L1 | miR-17-5p/20-5p/93-5p/106-5p/519-3p | + |

**Supplementary Table 6.** List of regulatory connections from Supplementary Table 2 that were labeled as high-confidence candidates with pan-cancer relevance (also illustrated in Fig. 4f). The Type column represents the type of interaction, where a plus (+) sign represents an activating interaction and a minus (-) sign represents an inhibitory interaction.

**A**

| miRNA | Target | Interaction (pair) type | Forward primer | Reverse primer | RT primer | Validated |
|---|---|---|---|---|---|---|
| miR-34-5p/449-5p | RRP1B | Predicted | GGGTGGCAGTGTCTTAGCTG | TTGAGAGGAAAGGTCGCTGAG | CCGCATTTGAGAGGAAAGGTC | Yes |
| miR-758-3p | MAP2K1 | Predicted | CGTTTGTGACCTGGTCCAC | TGGTACTTGGATATGGGGAAAG | CAGCATTGGTACTTGGATATGGG | Yes |
| miR-376c-3p | ACSL1 | Predicted | CCCAACATAGAGGAAATTCCAC | GCCTCACTCACCTGAAATGC | AAAAGATGCCTCACTCACCTG | Yes |
| miR-376c-3p | FCHO2 | Predicted | CCCAACATAGAGGAAATTCCAC | AGGCCTGATATTACACTATAACCC | TTTGTGCTTGCTCTTCTGAAAC | Yes |
| miR-10-5p | HOXA1 | Predicted | GTACCCTGTAGATCCGAATTTG | GTAAGAACAAATCAAGGCTACCC | CTCTGCAGGGCTTCTTAACTG | Yes |
| miR-10-5p | USF2 | Predicted | GTACCCTGTAGATCCGAATTTG | GGACCCAGAAACAAGAGGTG | GAAAATTAAAACACACACACACACC | No |
| miR-133a-3p.2/133b | UBA2 | Predicted | TATTTGGTCCCCTTCAACCAG | ACTGTGCAGGCATGTGTAAC | TATAAGGCAGCAGAAACAGGAAC | No |
| miR-377-3p | SESN2 | Predicted | AAACGATCACACAAAGGCAAC | TCCATTCCAGTGAGGAAAAGG | TAGGATCTCCAAGTGCTCTTCC | No |
| miR-377-3p | TMX3 | Predicted | AAACGATCACACAAAGGCAAC | ATATGAGGGAAACGGATGGC | AAATATATGAGGGAAACGGATGG | No |
| miR-182-5p | RARG | Predicted | GATTTGGCAATGGTAGAACTCAC | AGCCGGTTAGATCAGGAAGG | GAGGGAGCCGGTTAGATCAG | Yes |
| miR-182-5p | ARF4 | Predicted | GATTTGGCAATGGTAGAACTCAC | CCAGGCATTTACGCTTATGGG | CTATGTCCAGGCATTTACGC | Yes |
| miR-27-3p | TRAF1 | Predicted | CCCTTCACAGTGGCTAAGTTC | CCAGCCTTCACCCATCTTTG | AGTCGACCCCTCAGTCTTG | No |
| miR-29-3p | DSTYK | Predicted | CCCTAGCACCATTTGAAATCAG | CAACTCTTCACTGTCCAGGAG | TGCTCAGTCATTCAACTCTTCAC | No |
| miR-29-3p | C5orf15 | Predicted | CCCTAGCACCATTTGAAATCAG | TGGCGGAGAATGAATGTGAG | GTATTTGAATGGCGGAGAATG | No |
| miR-34-5p/449-5p | USF2 | Negative control | GGGTGGCAGTGTCTTAGCTG | GGACCCAGAAACAAGAGGTG | GAAAATTAAAACACACACACACACC | No |
| miR-758-3p | ACSL1 | Negative control | CGTTTGTGACCTGGTCCAC | GCCTCACTCACCTGAAATGC | AAAAGATGCCTCACTCACCTG | No |
| miR-376c-3p | PAM | Negative control | CCCAACATAGAGGAAATTCCAC | TCCAGGCGATTCTACTGTGG | AATTCTCCAGGCGATTCTACTGTG | No |
| miR-10-5p | UBA2 | Negative control | GTACCCTGTAGATCCGAATTTG | ACTGTGCAGGCATGTGTAAC | TATAAGGCAGCAGAAACAGGAAC | No |
| miR-133a-3p.2/133b | TMX3 | Negative control | TATTTGGTCCCCTTCAACCAG | ATATGAGGGAAACGGATGGC | AAATATATGAGGGAAACGGATGG | No |
| miR-377-3p | MAP2K1 | Negative control | AAACGATCACACAAAGGCAAC | TGGTACTTGGATATGGGGAAAG | CAGCATTGGTACTTGGATATGGG | No |
| miR-182-5p | TRAF1 | Negative control | GATTTGGCAATGGTAGAACTCAC | CCAGCCTTCACCCATCTTTG | AGTCGACCCCTCAGTCTTG | No |
| miR-27-3p | DSTYK | Negative control | CCCTTCACAGTGGCTAAGTTC | CAACTCTTCACTGTCCAGGAG | TGCTCAGTCATTCAACTCTTCAC | No |
| miR-29-3p | RARG | Negative control | CCCTAGCACCATTTGAAATCAG | AGCCGGTTAGATCAGGAAGG | GAGGGAGCCGGTTAGATCAG | No |
| miR-34-5p/449-5p | SIRT1 | Positive control | GGGTGGCAGTGTCTTAGCTG | CATCTATAGCACACAAACATCATGC | GGAAGATATCCAGTCATTAAACAGTC | Yes |

**B**

| | Predicted miRNA-target pairs | Negative control pairs |
|---|---|---|
| Validated | 7 | 0 |
| Not Validated | 7 | 9 |

**Supplementary Figure 1. Validation of predicted miRNA targets using AGO2-CLASH. (a)** Sequences of qRT-PCR primers designed to capture miRNA-mRNA hybrids. **(b)** Table showing the number of miRNA-mRNA pairs that were validated by AGO2-CLASH, among predicted targets and negative control pairs. 14 predicted miRNA-mRNA interactions from our network were tested, encompassing 9 different miRNAs. As negative controls, 9 random pairs were included, encompassing the same miRNAs matched to random mRNAs that are not predicted targets.

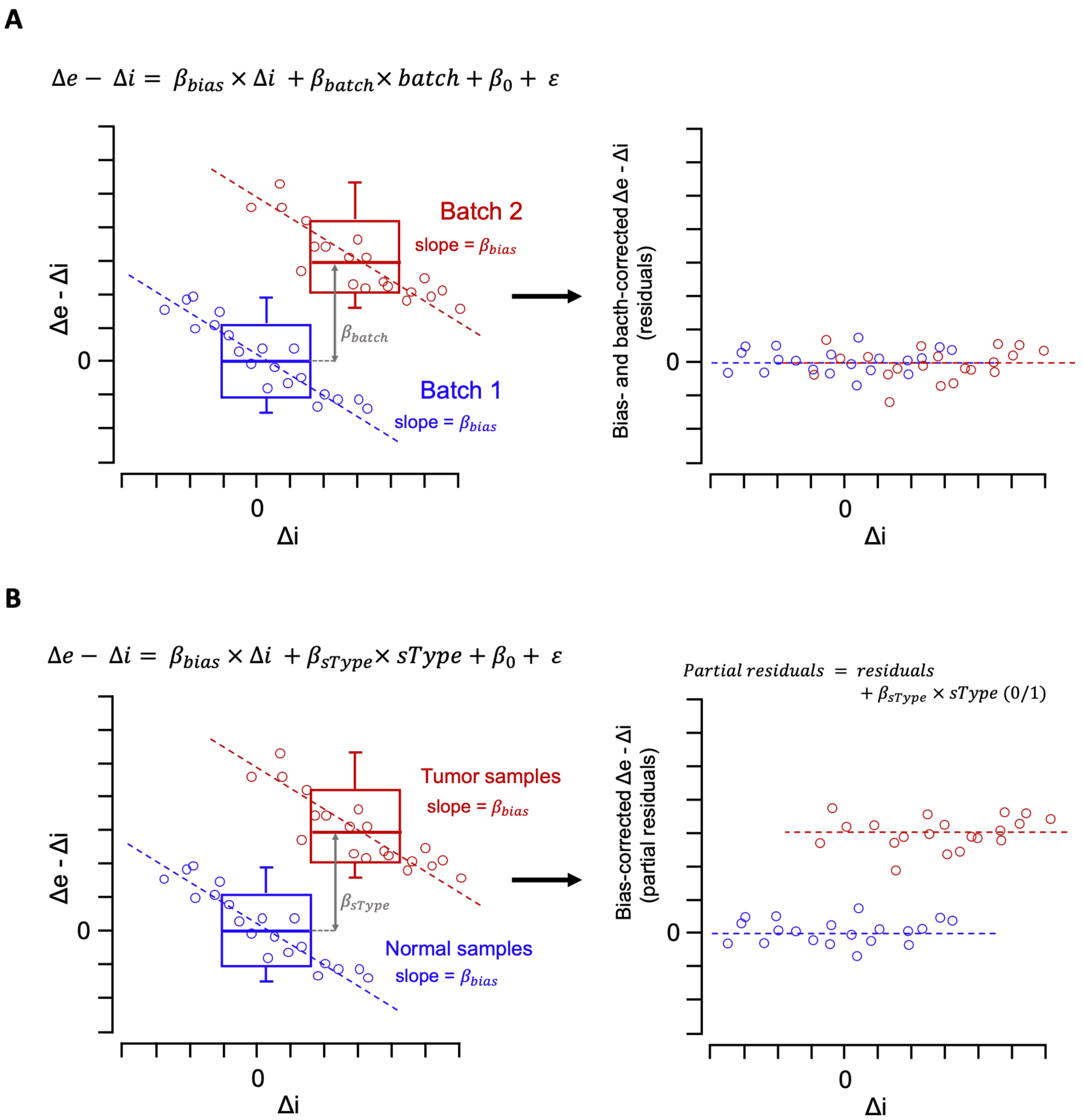


**Supplementary Figure 2. Correction of mRNA stability estimates for transcription-associated bias and for experimental batch.** (**a**) Schematic illustration of the model used to correct mRNA stability estimates in ENCODE RBP knockdown samples for the confounding effects of the batch and transcription-associated bias (see Methods). The equation on top represents the model used, with the interpretation of model coefficients shown on the plot. The data points and box plots are drawn manually for illustration purposes. (**b**) Similar to (a), for mRNA stability estimates from TCGA samples.

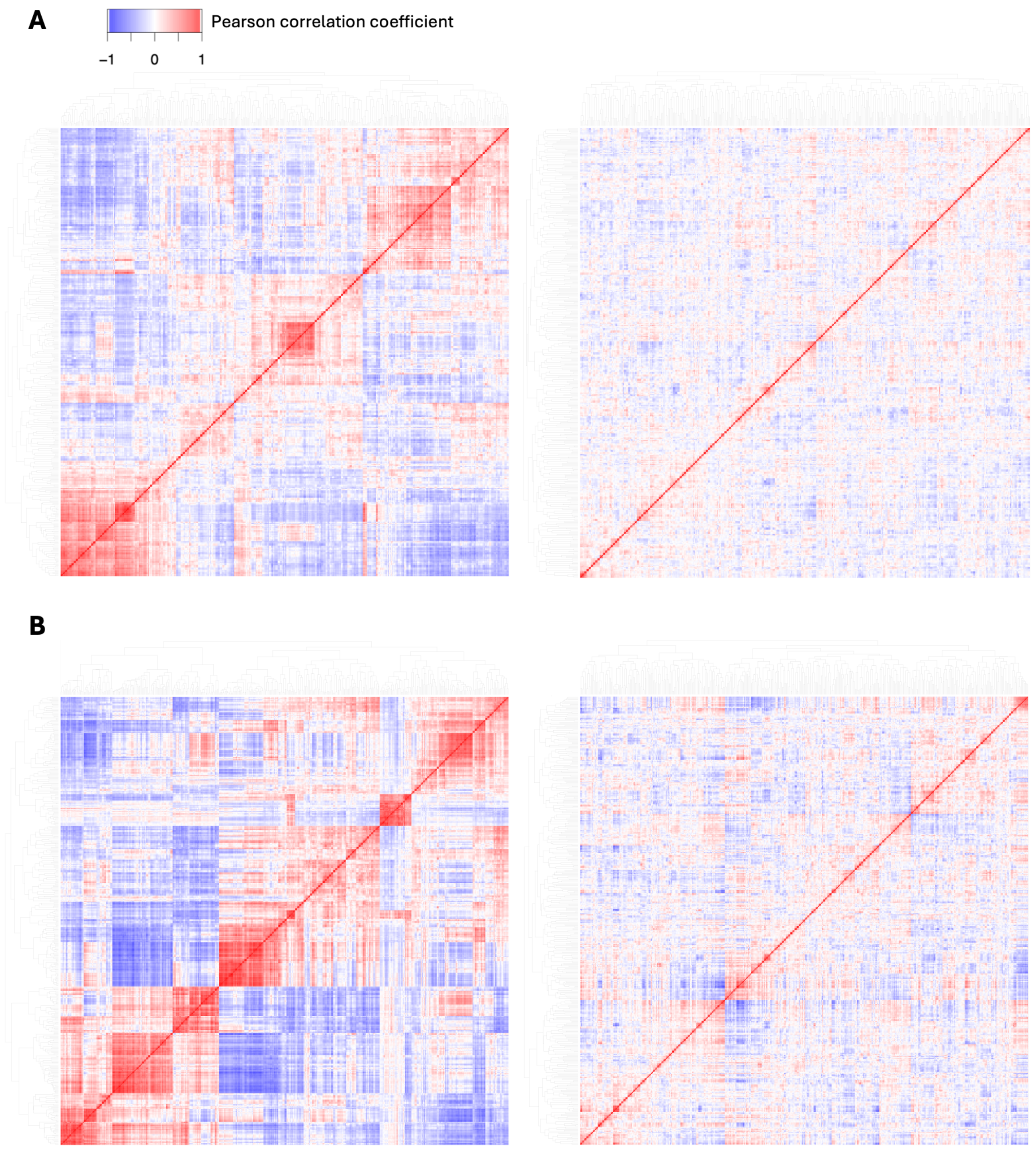


**Supplementary Figure 3. Batch correction of mRNA stability estimates in RBP knockdown experiments from ENCODE.** (**a**) Correlation heatmaps of mRNA stability estimates in RBP knockdown experiments from ENCODE, in HepG2 cells before (left) and after (right) batch correction (see Methods). The color gradient represents the Pearson correlation coefficient for pairwise comparison of samples. Each row and column represent one RBP knockdown sample. (**b**) Similar to panel (a), for RBP knockdowns in K562 cells.

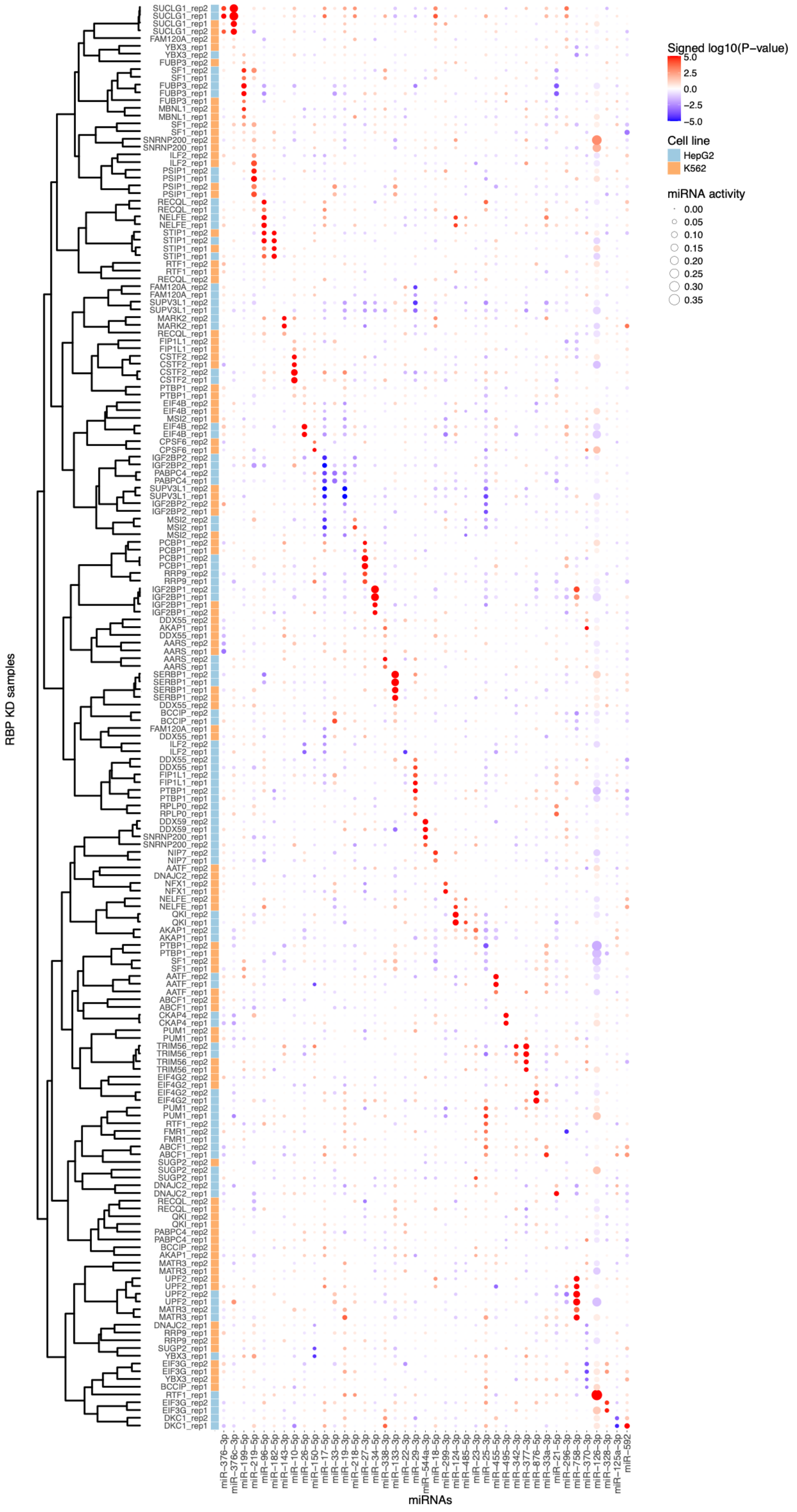


**Supplementary Figure 4. miRNA activity profiles in RBP knockdown experiments before correction for shRNA off-target effects.** Similar to Fig. 2a, before correction for shRNA off-target effects.

**A**

```
shIGF2BP1       CCGGGCAGTGGTGAATGTCACCTATCTCGAGATAGGTGACATTCACCACTGCTTTTTG
miR-34 seed        GGCAGUG
```

**B**

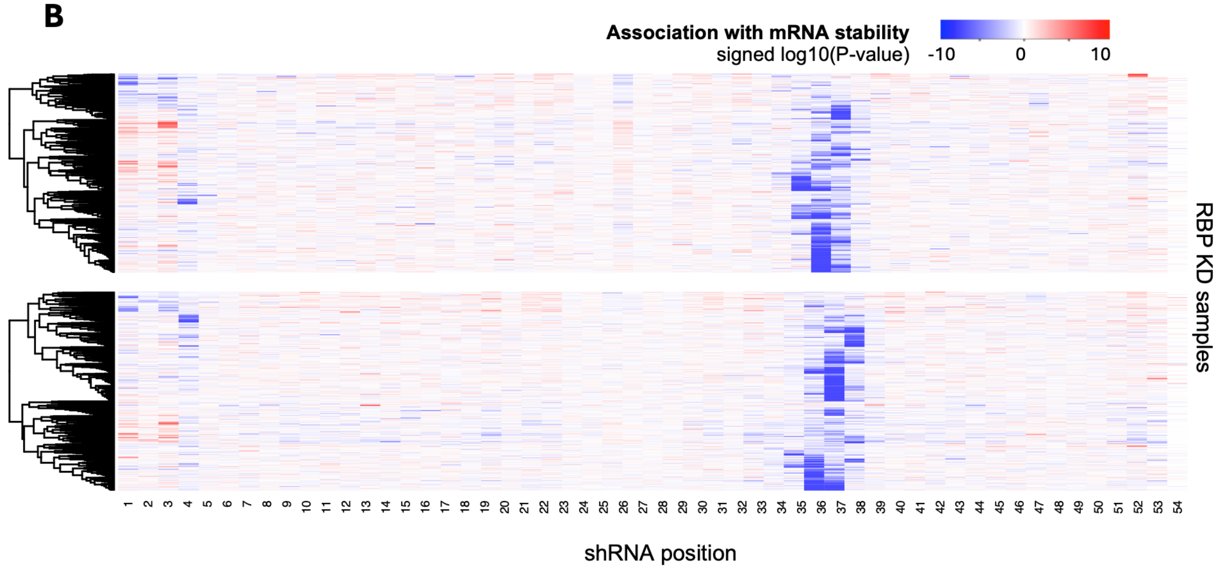


**Supplementary Figure 5. Specific shRNA positions mediate off-target effects on mRNA stability.** (**a**) An example shRNA containing a miRNA seed sequence, for which a negative RBP-miRNA interaction was initially inferred (see Supplementary Fig. 4). (**b**) shRNA positions associated with off-target effects on mRNA stability (see Methods). The color gradient represents the signed log10(P-value) of the association between counts of each shRNA 6-mers (top) or 7-mers (bottom) in each gene's 3'UTR and changes in mRNA stability, obtained using a linear model. Each row represents one RBP knockdown sample (and therefore one shRNA), and each column represents the nucleotide positions on the shRNA. Blue indicates a negative effect on mRNA stability and red indicates a positive effect.

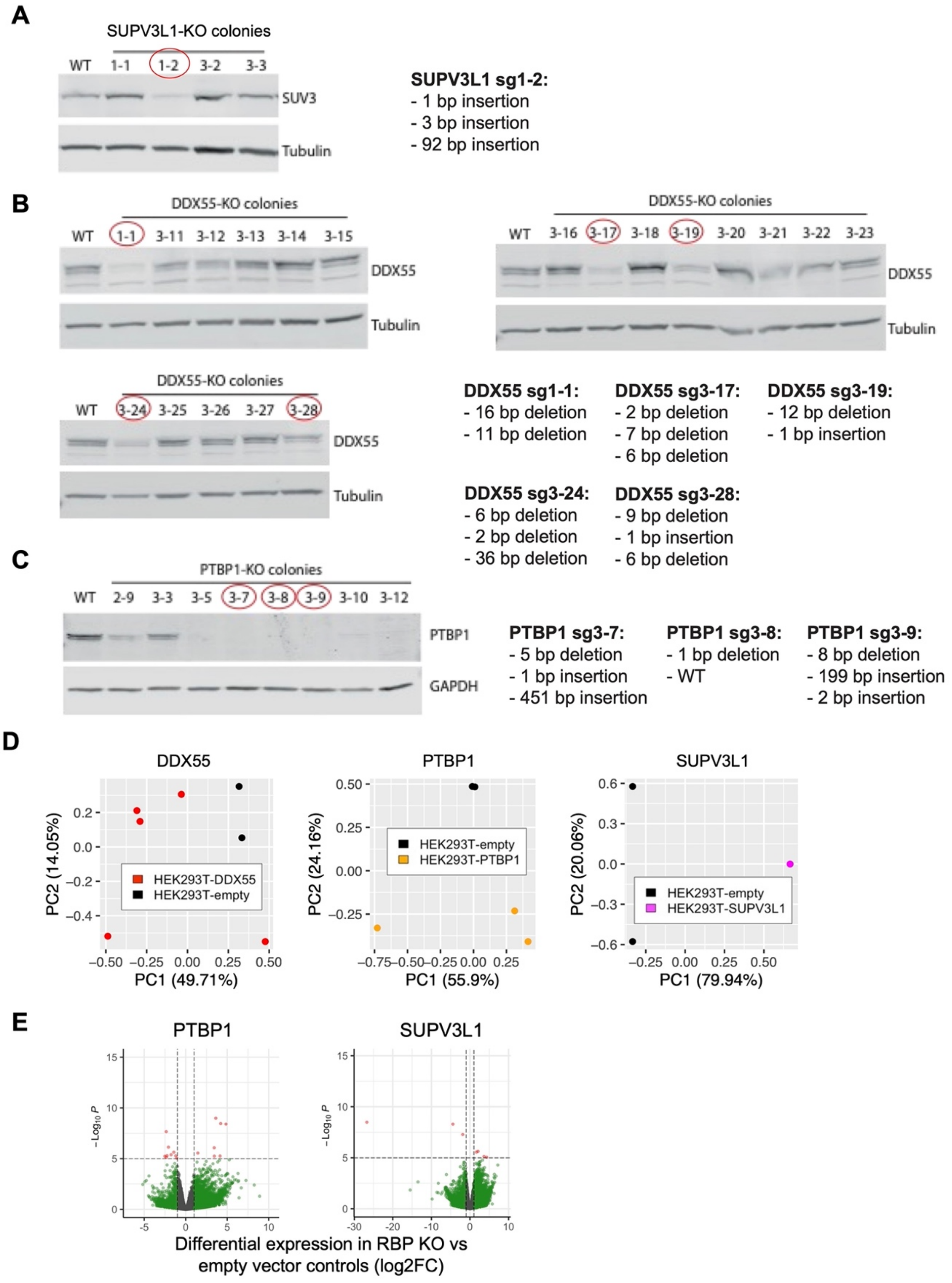


**Supplementary Figure 6. Validation of CRISPR/Cas9-mediated knockout cell lines.** Western blot analysis of (**a**) PTBP1 (**b**) DDX55 and (**c**) SUPV3L1 protein expression in HEK293T wild-type (WT)

and knockout clones. GAPDH expression was used as a loading control. Mutations at each allele for each clone are indicated beside the blots. Clones 3-8 and 3-9 showed complete loss of PTBP1 (a) and are circled in red. For DDX55 inhibition (b), no clones with complete loss of DDX55 were viable. 5 clones (circled in red) showed reduced DDX55 expression. Similarly, for SUPV3L1 inhibition (c), no clones with complete loss of SUPV3L1 were viable. A single clone (circled in red) showed reduced SUPV3L1 expression and was retained. (**d**) Principal component analysis (PCA) of RNA-seq read counts in knockout samples following variance-stabilized transformation from DESeq2 and centering each gene by subtracting its mean across all samples. (**e**) Volcano plots of differential gene expression in RBP knockouts compared to empty vector controls.

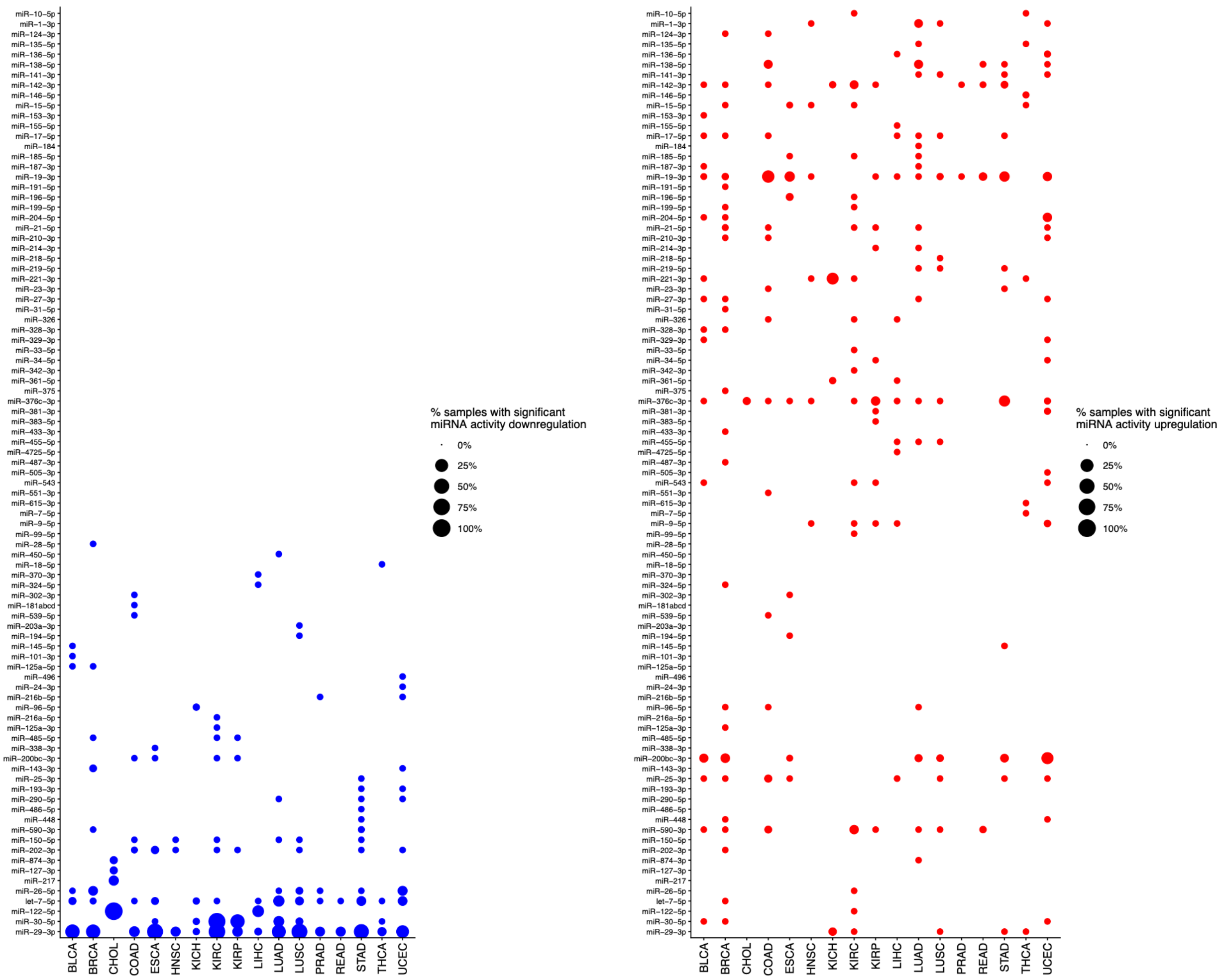


**Supplementary Figure 7. Differential miRNA activity in TCGA cancers.** Heatmap showing the differentially active miRNAs in each cancer types. The circle size represents the percent of tumor samples with a significant (FDR < 0.05) increase (right, red) or decrease (left, blue) change in miRNA activity relative to the average of normal samples, within each cancer type. miRNAs with a significant change in activity in either direction in at least one tumor are shown.

A

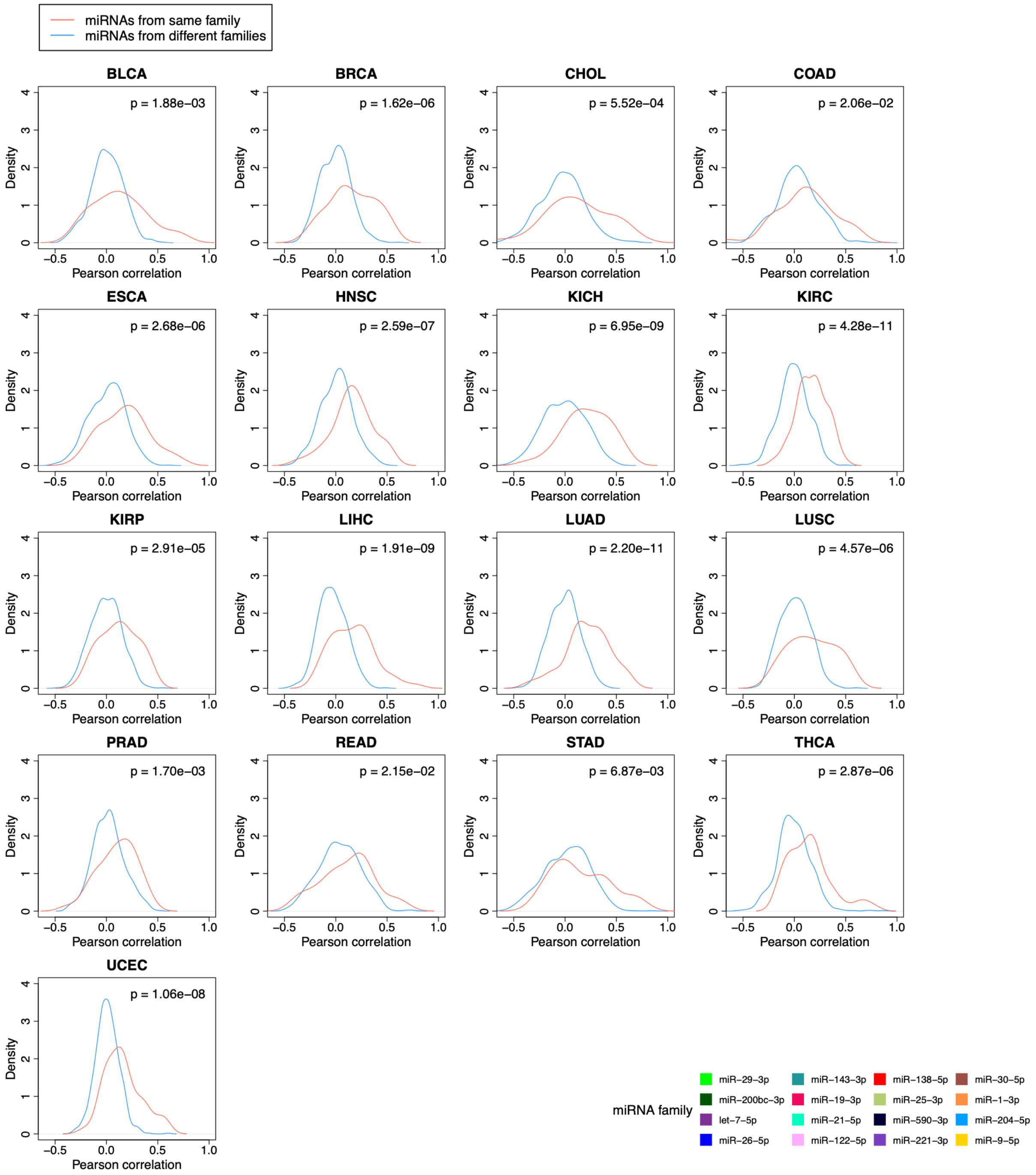

miRNAs from same family
miRNAs from different families
BLCA
p = 1.88e−03
BRCA
p = 1.62e−06
CHOL
p = 5.52e−04
COAD
p = 2.06e−02
ESCA
p = 2.68e−06
HNSC
p = 2.59e−07
KICH
p = 6.95e−09
KIRC
p = 4.28e−11
KIRP
p = 2.91e−05
LIHC
p = 1.91e−09
LUAD
p = 2.20e−11
LUSC
p = 4.57e−06
PRAD
p = 1.70e−03
READ
p = 2.15e−02
STAD
p = 6.87e−03
THCA
p = 2.87e−06
UCEC
p = 1.06e−08
Density
Pearson correlation
miRNA family
miR−29−3p
miR−200bc−3p
let−7−5p
miR−26−5p
miR−143−3p
miR−19−3p
miR−21−5p
miR−122−5p
miR−138−5p
miR−25−3p
miR−590−3p
miR−221−3p
miR−30−5p
miR−1−3p
miR−204−5p
miR−9−5p


B

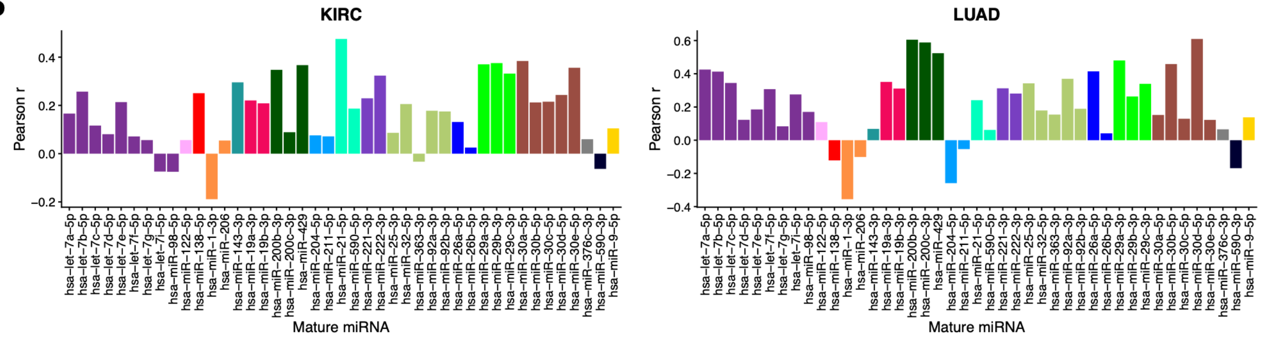

KIRC
LUAD
Pearson r
Mature miRNA

**Supplementary Figure 8. Agreement between inferred miRNA activity and expression.** (**a**) Probability density functions of Pearson correlations between inferred activity and expression for miRNAs of the same family (red), compared to miRNAs from different families (blue), in all TCGA cancers for which activity was inferred. miRNAs with significant changes in activity in at least 10 tumor samples in at least one cancer type are included (see Supplementary Table 4). The P-values from the Mann-Whitney U test comparing the two distributions are shown on the plots. (**b**) Correlation of inferred activity of each miRNA family included in (a) with the expression of each of their family members, in TCGA-KIRC (left) and TCGA-LUAD (Lung adenocarcinoma) (right), shown as examples. The colors represent each miRNA family, and each bar represents one family member.

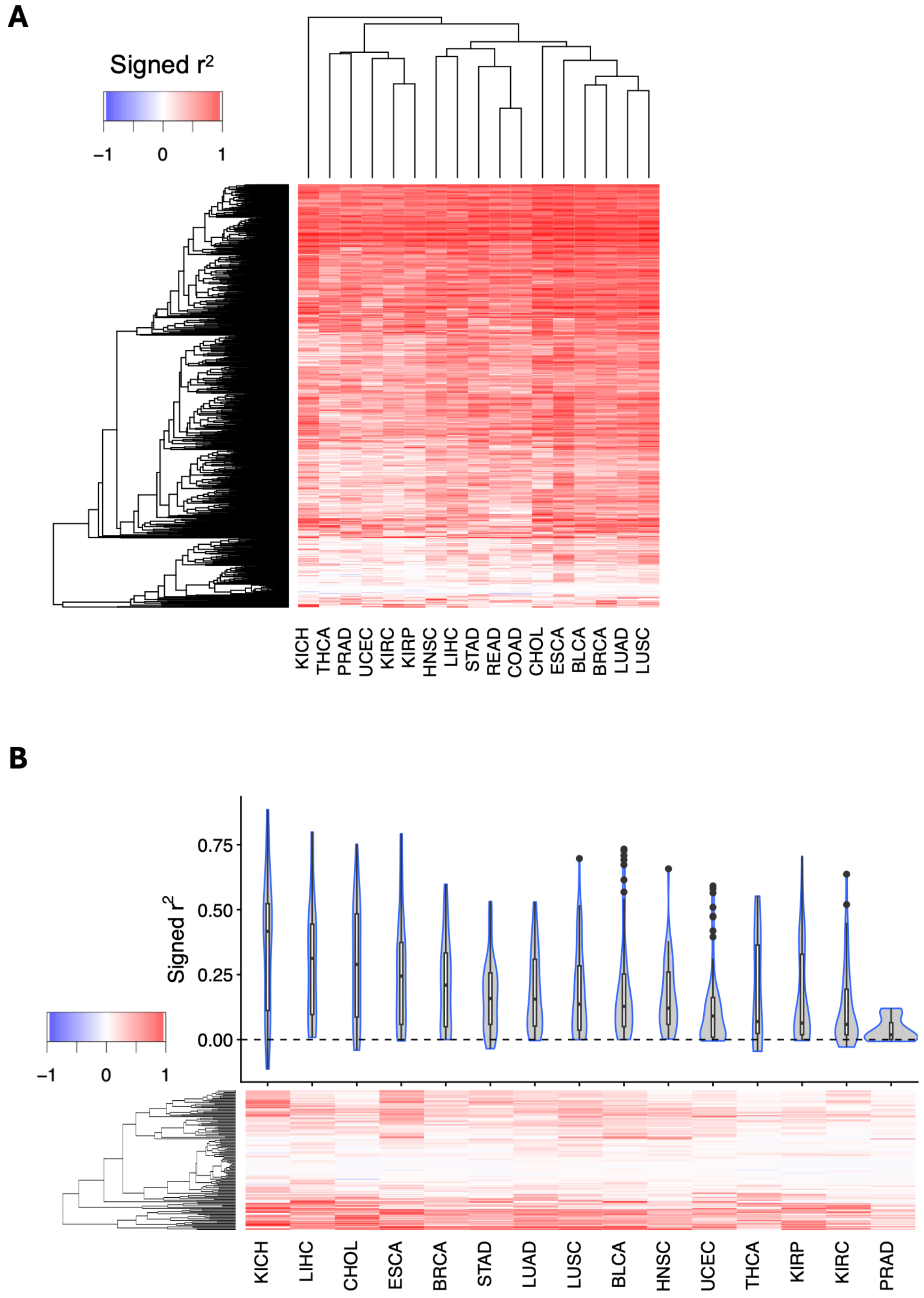


**Supplementary Figure 9. Association between miRNA transcription and abundance across TCGA cancers.** (**a**) Association between miRNA host gene transcription and abundance. Transcriptional changes were inferred from changes in intronic read counts, while changes in abundance were inferred from changes in exonic read counts, both relative to the average. Associations were tested using a linear model and signed $r^2$ values ($r^2$ values with the sign corresponding to that of the Pearson correlation coefficient) are shown. (**b**) Association between miRNA transcription and mature miRNA abundance in TCGA tumors, for differentially expressed miRNAs. Differentially expressed miRNAs were identified from mature miRNA abundance data log2 (RPM) from small RNA-seq, using matched tumor-normal sample pairs from the same patient. A paired t-test was used and miRNAs with $|log2FC| > 1$ and adjusted P-value $< 0.05$ were retained. Transcriptional changes were inferred from that of the host genes, which were based on changes in intronic read counts (see Methods). Both the violin plots (top) and heatmap (bottom) show the signed $r^2$ for each cancer type. Each data point in the violin plots and each row in the heatmap represent one miRNA. The heatmap includes all miRNAs that were significant in at least one tumor type.

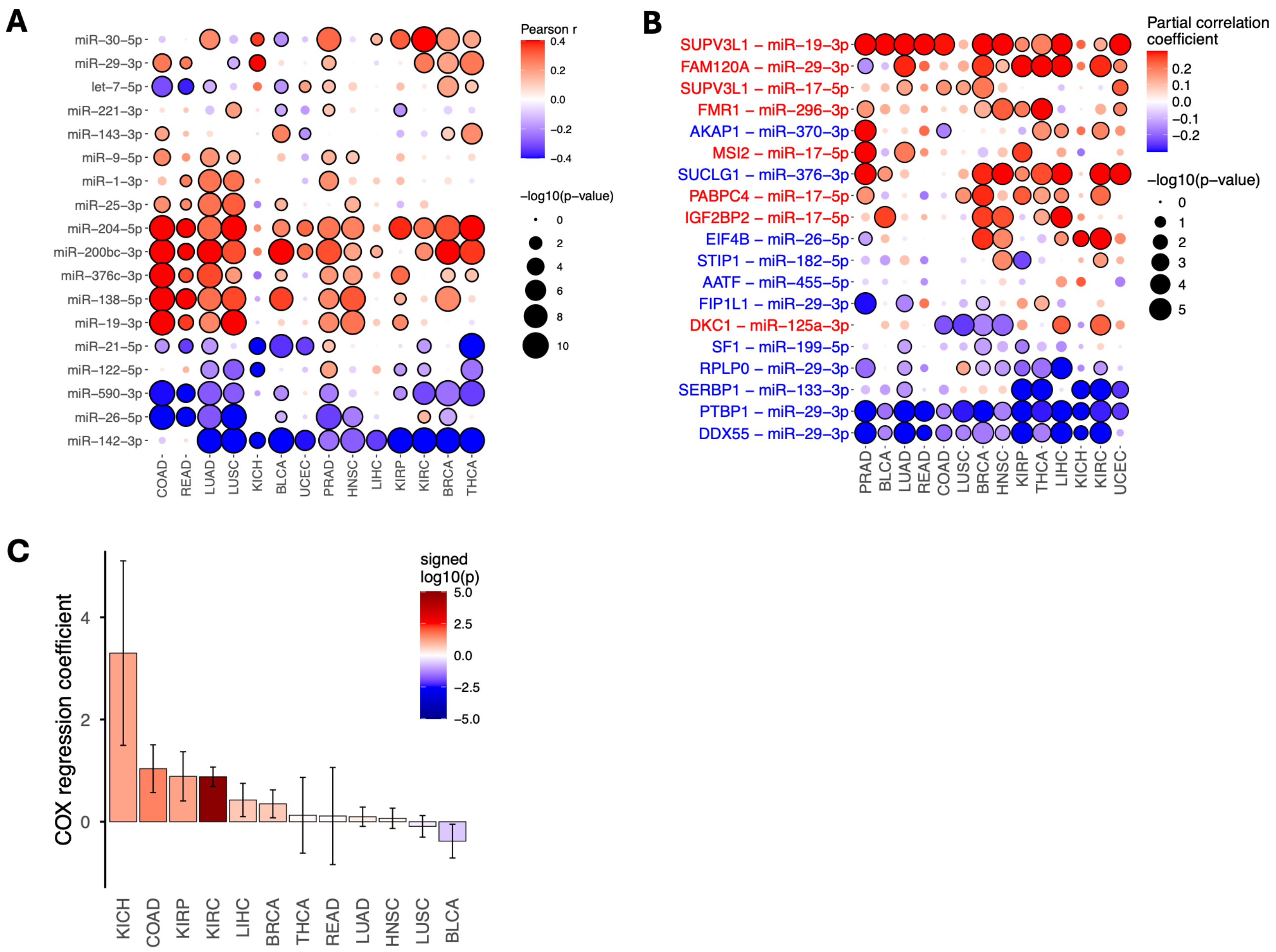


**Supplementary Figure 10. Evaluating the impact of tumor purity on TCGA data analyses.** (**a**) Correlation between inferred miRNA activity and tumor purity, for miRNAs with significant changes in activity in at least 10 tumor samples in at least one cancer type (see Supplementary Table 4). The color gradient represents the correlation coefficients; the circle sizes represent the -log10(P-values) and a black outline indicates a significant result at FDR < 0.05. (**b**) Similar to Fig. 4b, correlations between miRNA activity and RBP expression across TCGA cancers. Here, partial correlations (i.e. the correlation of miRNA activity and RBP expression while controlling for tumor purity) are computed, using the *pcor.test* function in R [1]. (**c**) Similar to Fig. 5c, associations between DDX55 expression and survival, with added correction for patient age, tumor stage and purity.

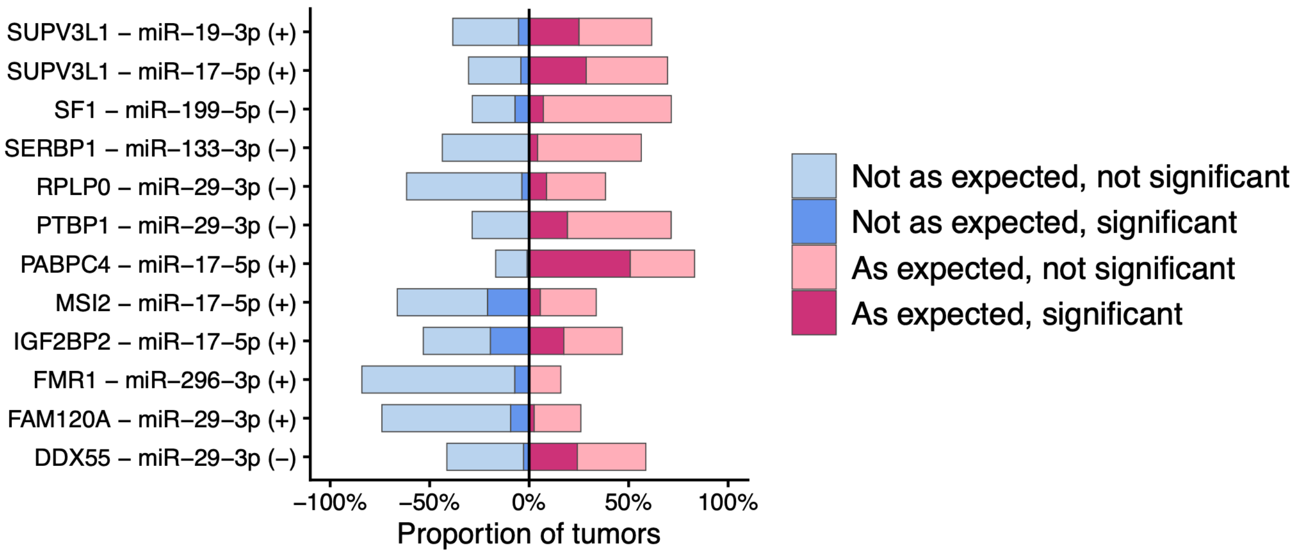


**Supplementary Figure 11. Behavior of RBP-responsive miRNA targets in cancer.** Diverging stacked bar chart indicating the percent of TCGA tumor samples that show enrichment of the RBP-KD-consistent miRNA targets, among differentially stable genes in each TCGA tumor samples. See Fig. 4c for more detail.

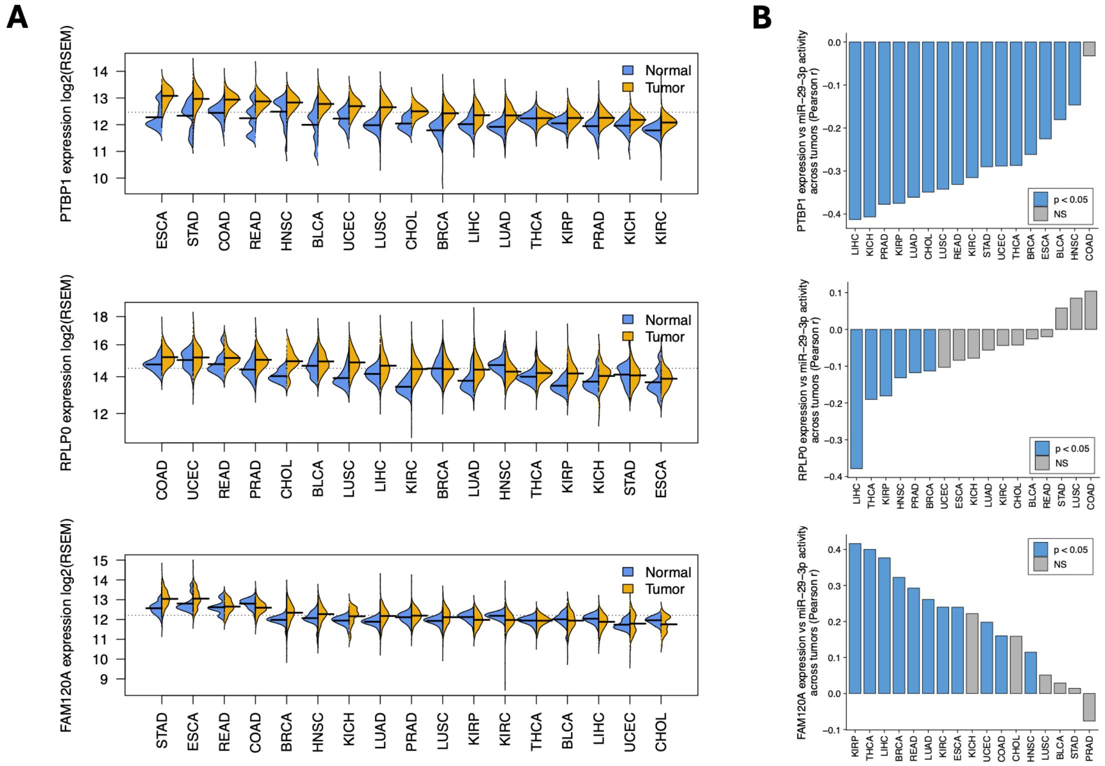


**Supplementary Figure 12. Expression of other candidate RBPs interacting with miR-29-3p in cancer and their association with miRNA activity.** Comparison of 3 RBPs potentially interacting with miR-29-3p other than DDX55 (PTBP1, RPLP0 and FAM120A) in terms of relevance to cancer. (**a**) Expression across TCGA datasets, shown as log2(RSEM). (**b)** Correlations between RBP expression and miRNA activity across TCGA datasets. Blue represents significant associations (also see Fig. 4b).

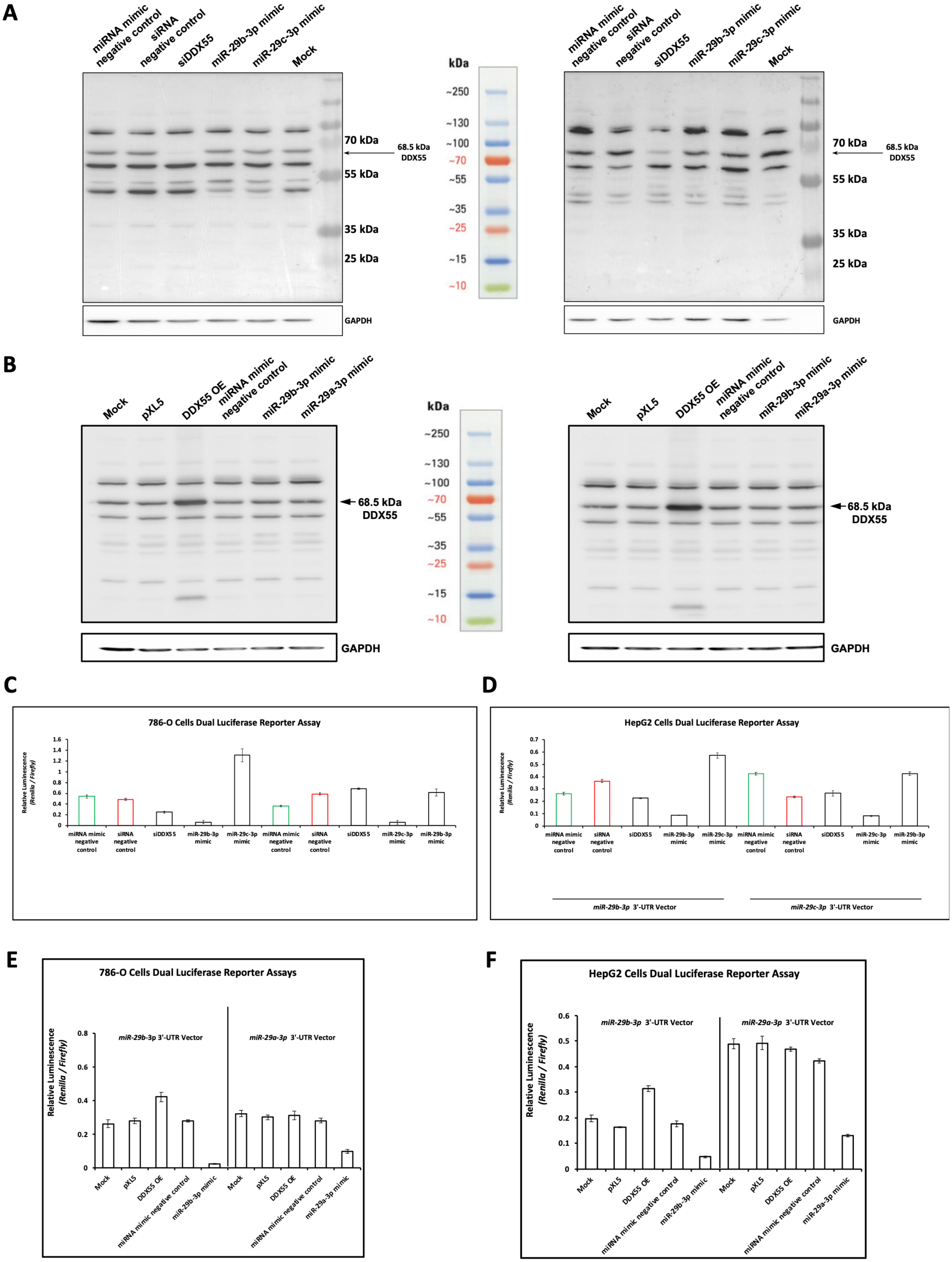
A
miRNA mimic negative control
siRNA negative control
siDDX55
miR-29b-3p mimic
miR-29c-3p mimic
Mock
70 kDa
55 kDa
35 kDa
25 kDa
68.5 kDa DDX55
GAPDH
kDa
~250
~130
~100
~70
~55
~35
~25
~15
~10
B
Mock
pXL5
DDX55 OE
miRNA mimic negative control
miR-29b-3p mimic
miR-29a-3p mimic
68.5 kDa DDX55
GAPDH
C
786-O Cells Dual Luciferase Reporter Assay
Relative Luminescence (Renilla / Firefly)
miRNA mimic negative control
siRNA negative control
siDDX55
miR-29b-3p mimic
miR-29c-3p mimic
D
HepG2 Cells Dual Luciferase Reporter Assay
miR-29b-3p 3'-UTR Vector
miR-29c-3p 3'-UTR Vector
E
786-O Cells Dual Luciferase Reporter Assays
miR-29b-3p 3'-UTR Vector
miR-29a-3p 3'-UTR Vector
Relative Luminescence (Renilla / Firefly)
Mock
pXL5
DDX55 OE
miRNA mimic negative control
miR-29b-3p mimic
miR-29a-3p mimic
F
HepG2 Cells Dual Luciferase Reporter Assay
miR-29b-3p 3'-UTR Vector
miR-29a-3p 3'-UTR Vector

**Supplementary Figure 13. Validation of DDX55 knockdown and DDX55 overexpression in cell lines and their impact on miR-29 activity.** Western blot analyses confirming the (**a**) siRNA-mediated knockdown and (**b**) overexpression of DDX55 in 786-O (left) and HepG2 (right) cell lines. GAPDH expression was used as a loading control. (**c**) Relative luciferase luminescence (Renilla vs Firefly), representing the inverse of miR-29b-3p (left) or miR-29c-3p (right) activity, upon siRNA-mediated DDX55 knockdown in 786-O cells and subsequent expression of miR-29b-3p mimic, miR-29c-3p mimic, siRNA negative control, or miRNA mimic negative control. Error bars represent SD, for 5 biological replicates. (**d**) Similar to (c), in HepG2 cells. (**e**) Relative luciferase luminescence (Renilla vs Firefly), representing the inverse of miR-29b-3p (left) and miR-29a-3p (right) activity, upon DDX55 overexpression in 786-O cells and subsequent expression of a miR-29b-3p mimic, miR-29a-3p mimic, empty vector control (pXL5), or miRNA mimic negative control. Error bars represent SD, for 5 biological replicates. (**f**) Similar to (e), in HepG2 cells.

### References

1 Kim, S. ppcor: An R Package for a Fast Calculation to Semi-partial Correlation Coefficients. *Commun Stat Appl Methods* **22**, 665-674, doi:10.5351/CSAM.2015.22.6.665 (2015).
